# Early-life stress impairs development of functional interactions and neuronal activity within prefrontal-amygdala networks *in vivo*

**DOI:** 10.1101/2024.12.04.626305

**Authors:** Angelica Donati, Francescangelo Vedele, Henrike Hartung

**Affiliations:** HiLIFE Neuroscience Center, P.O. 63, FI-00014 University of Helsinki, Helsinki, Finland; Department of Physiology, Medicum, University of Helsinki, Helsinki, Finland

## Abstract

Early-life stress (ELS), such as parental neglect or abuse, predisposes an individual to develop mental disorders. Disease hallmarks include heightened amygdala reactivity and impaired prefrontal cortex-amygdala functional interactions, already during childhood and adolescence. However, which cellular and circuit mechanisms underlie these hallmarks, as well as the altered developmental trajectory of prefrontal-amygdala networks, is poorly understood. Here we performed simultaneous *in vivo* local-field potential and multi-unit recordings under light urethane anaesthesia in the medial prefrontal cortex (mPFC) and basolateral amygdala (BLA) of male and female pre-juvenile or adolescent mice, exposed to a resource scarcity model of ELS. We find a developmentally transient low-theta (3-5 Hz) oscillatory hypercoupling within mPFC-BLA networks in pre-juvenile ELS males which seems to result from a precocious development of coupling strength after ELS. In the mPFC, neuronal spiking activity was decreased in pre-juvenile males and the local theta entrainment of spike firing disrupted. In BLA, both sexes showed an increase in firing activity in a subpopulation of neurons after ELS, also confirmed by an increase in ΔFosB-positive neurons in BLA, which we identified to be non-GABAergic. Directed interactions, i.e. the ability to entrain spike firing in mPFC to the theta rhythm in BLA and vice versa, were also impaired predominantly in pre-juvenile males after ELS, while females showed a milder phenotype. These early sex-dependent impairments in the functional development of prefrontal-amygdala circuits may promote aberrant development of emotional behaviours after ELS and may predispose to a disease phenotype later on.

## 1. Introduction

Early-life stress (ELS) such as parental neglect, maltreatment, or abuse, is a strong risk factor for depression and anxiety disorders ^1–3^. Studies in humans and animal models have pointed to malfunction of prefrontal-amygdala networks as the substrate increasing the risk for psychiatric symptoms after the experience of ELS ^4, 5^. Both the amygdala and the prefrontal cortex are particularly vulnerable to early stressors due to their protracted postnatal development. Most extensively for the amygdala, but also for other limbic brain regions, it has been shown that ELS leads to an accelerated structural and functional maturation across species, i.e. a precocious development of emotion circuits ^6^. The most consistent findings among individuals with a history of ELS are amygdala hyper-reactivity to (mainly) negative emotional cues as well as impaired prefrontal cortex-amygdala task-elicited or resting-state functional connectivity detected as early as childhood ^4, 5^. How these ELS hallmarks emerge and determine each other, particularly concerning the protracted and sequential development of this circuit, is still hardly understood.

The recently developed limited bedding and nesting (LBN) model mimics resource scarcity and altered parental care, which is often related to early adversity in humans ^7, 8^. The altered rearing condition induces stress to the dam and leads to unpredictable maternal care patterns ^7–9^. Mice reared under LBN conditions display delayed physical growth and deficits in spatial ^10, 11^ and contextual learning ^12, 13^ as well as depressive-like behaviours ^14^ and increased anxiety ^15^ in a sex-dependent manner. Furthermore, hyper-reactivity of amygdala networks, as measured by a stimulus-induced increase in the immediate early gene cFos-positive neurons in the basolateral amygdala (BLA), has also been found in this model, ^17^. *In vitro* electrophysiological recordings after ELS showed increased excitatory input to BLA neurons and increased synaptic plasticity (LTP) ^18, 19^ predominantly in males, but also an increase in the intrinsic excitability of BLA neurons likely mediated by reduced expression of SK channels ^20^. However, how ELS affects neuronal spiking activity within prefrontal-amygdala networks *in vivo* is still largely unknown. In line with data in humans, ELS rodent models also show impaired prefrontal-amygdala functional connectivity as measured by fMRI or tractography predominantly in males, that strongly correlate with behavioural impairments in innate anxiety or fear conditioning ^15, 21–26^. However, direct electrophysiological measurements of how ELS affects the development of local prefrontal or amygdala network activity and their long-range interactions remain sparse.

Here, we used a cumulative combination of the LBN resource scarcity model and maternal separation (MS) to investigate the effect of ELS on *in vivo* network activity, as well as long-range functional interactions between prelimbic (PL) or infralimbic (IL) subdivisions of the medial prefrontal cortex (mPFC) and BLA during pre-juvenile (postnatal day (P) 18-20) or adolescent (P43-47) development in male and female pups. Cumulative ELS exposure to LBN+MS has previously been shown to lead to an anxiety phenotype in male mice as well as aberrant prefrontal-amygdala functional and structural connectivity ^15, 24^. Besides, initial behavioural characterization of our cumulative ELS model during adulthood also revealed increased anxiety-like behaviours in ELS males and strong deficits in classical fear conditioning in both ELS males and females^27^. In this study, we found functional impairments within prefrontal-amygdala networks after ELS already during pre-juvenile development predominantly in males, such as oscillatory hypercoupling in the low-theta band (3-5 Hz) as well as disrupted directed interactions within prelimbic-BLA networks. Moreover, in the mPFC neuronal spiking activity was decreased, and local theta entrainment was disrupted in ELS males. In contrast, in pre-juvenile BLA, a subpopulation of presumably principal neurons showed increased firing activity after ELS in both sexes. This effect was even more prominent during adolescent development with a broad increase in mean firing rates in ELS males. This study is one of the first investigating the impact of ELS on the functional development of prefrontal-amygdala networks by direct *in vivo* electrophysiological measurements. Our data provide further evidence for pre-juvenile development as a susceptible period following ELS, during which both an impairment in functional long-range interactions as well as an abnormal local neuronal firing activity emerge in the PL and in the BLA. These early impairments may lead to major circuit dysfunction underlying disease in adulthood.

## 2.2 Materials and Methods

### 2.1 Animals

Experiments were performed using male and female C57Bl/6J mice. All experiments were done in accordance with the University of Helsinki Animal Welfare Guidelines and approved by the Animal Experiment Board in Finland (license ESAVI/13422/2018 and ESAVI/27422/2022). Mice were maintained on a 12-hour light-dark cycle (lights on from 6 am to 6 pm) with *ad libitum* access to food and water. Mice were housed in a quiet room adjacent to the laboratory to minimize stress from transportation.

### 2.2 Early-life stress protocol (limited bedding and nesting + maternal separation)

The limited bedding and nesting (LBN) protocol was a modified version of the protocol used by KG Bath and co-workers, see ^28^. In total 48 litters between 4-8 pups including at least two pups of each sex were randomly assigned to either control or LBN group. The average litter size was similar in both experimental groups (control: 6.0 ± 1.2 pups/litter, LBN: 6.2 ± 1.3 pups/litter). At P4, the dam and the entire litter were transferred to a cage with a wire mesh floor (L30cm x W30cm x H1cm) 1 cm above the cage floor and half a nestlet (2,3 x 5 cm) as the only source of nesting material. Food and water were supplied *ad libitum*. The LBN dam and litter remained in these housing conditions until P14. In this study, the LBN period was extended to include the developmental time window (P10-14) during which prefrontal cortical projections to BLA steeply increase ^29, 30^. In addition, pups were separated from their dams for 1 h on P8, P10, and P12 by moving the whole litter to a standard cage with normal bedding and nesting material, placed on a heating pad to maintain body temperature. Otherwise, the dam and pups were left undisturbed. Pups were weighed individually at P14 upon return to a standard home cage and were weaned at P21. Control litters were housed in standard home cages (L 42 cm x W 25 cm x H 15 cm) with a normal supply of bedding and nesting material as well as *ad libitum* access to food and water, and were left undisturbed between P4-14 and weaned at P21.

### 2.3 *In vivo* electrophysiology

#### Surgical preparation

Extracellular recordings were performed in the mPFC and in the anterior part of the BLA (BLAa) of P18-20 (pre-juvenile) and P43-47 (adolescent) male and female mice.

Pups were weighed and anaesthesia was induced with isoflurane (5%) followed by i.p. administration of urethane (1.2 g/kg; Sigma-Aldrich, USA). Isoflurane administration (1-2%) continued during surgery to ensure deep anaesthesia. An incision along the midline was made and the skull was exposed. Small burr holes were drilled above the regions of interest (mPFC: 1.2-1.3 mm anterior to bregma and 0.1 mm from the midline, BLA: 1.5-1.7 posterior to Bregma, at the base of the rhinal fissure) in the right hemisphere, and the bone was carefully removed to prevent tissue damage and bleeding. The head of the pup was fixed into the stereotaxic apparatus (RWD Life Science, China) using two hollow plastic bars fixed with dental cement on the nasal and occipital bones. The body temperature was maintained at 37 °C with a thermoregulated heating blanket (Supertech Instruments, UK). A urethane top-up of 1/5 of the original dose was administered if needed. Multielectrode arrays (32-channel, four-shank (4×8) silicon Michigan probes (0.4 -2 MΩ), A4x8-5mm-200-200-177-A32 (mPFC) or A4x8-5mm-100-200-177 (BLAa), NeuroNexus Technologies, USA) were inserted perpendicular to the skull surface into the mPFC until a depth of 1.8-2.0 mm, and at 40° from the vertical plane into the BLAa at a depth of 2.0-2.2 mm. The distance between shanks was 200 µm and the recording sites were separated by 100 µm (BLAa) or 200 µm (mPFC). The electrodes were labelled with DiI (1,1’-dioctadecyl-3,3,3’,3’-tetramethyl indocarbocyanine, Invitrogen, USA) to enable reconstruction of the electrode tracks postmortem (Figure 1B). A silver wire was inserted into the cerebellum and served as common ground and reference electrode.

**Figure 1.**
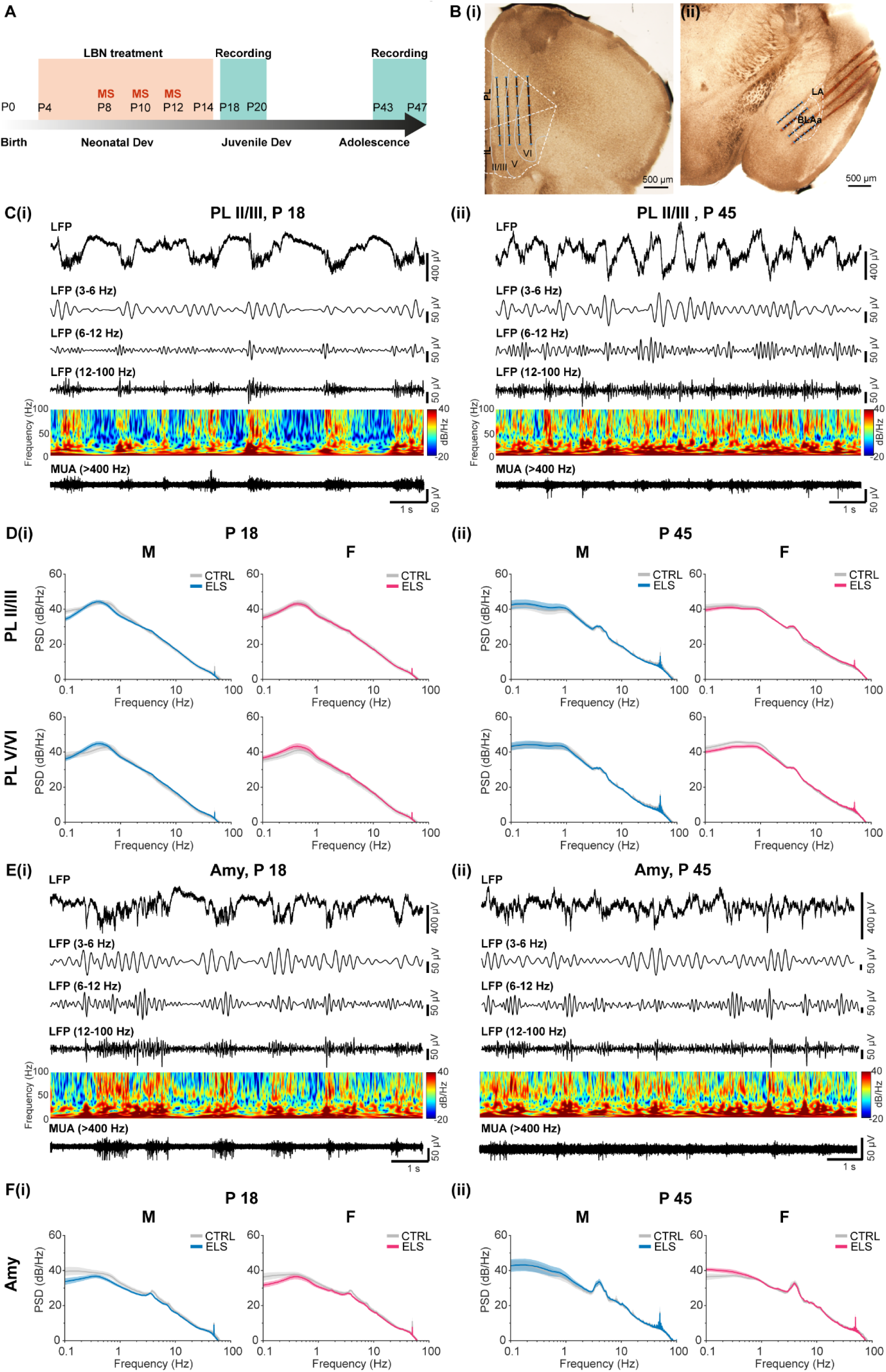
Oscillatory network activity in PL and BLAa of pre-juvenile and adolescent control mice. A, Diagram with timeline illustrating the experimental design. **B**, Digital photomontage of brightfield images with electrode tracts of the four-shank recording electrode and reconstruction of the 32 recording channels (blue dots) within **(i)** PL and IL or **(ii)** BLAa. **C, (i)** Extracellular LFP recording of oscillatory activity in PL, superficial layers II/III, of a male pre-juvenile mouse pup displayed after band pass filtering (3-6, 6-12, 12-100 Hz) and after 400 Hz high-pass filtering showing corresponding MUA activity. The color-coded frequency plot shows the wavelet spectra at identical time scale. Note the typical large-amplitude slow oscillations during urethane anaesthesia superimposed by theta, beta, and gamma band activity. Activity in the low-theta range (3-6 Hz) is particularly prominent. **(ii)** Same as C (i) for a male adolescent mouse pup. **D, (i)** Average logarithmic power spectra of prelimbic LFP in layers II/III (top) or layers V/VI (bottom) of male (left) and female (right) control (grey) or ELS (blue or magenta) pre-juvenile mice (males: N = 16-20/group, females: N = 13-18 /group). Note the broad peak in delta range (0.3-0.5 Hz) corresponding to slow-wave activity as well as the small peak in low-theta range at 4 Hz. **(ii)** Same as D (i) for adolescent control and ELS male and female mice (males: N = 13-16/group, females: N = 18-22/group). Note the shift of the peak in delta activity towards 1 Hz and the prominent broad peak in low theta activity (3-6 Hz). See Suppl. Table 2 for the exact number of pups per group in each graph in D. **E, (i)** Same as C (i) for corresponding oscillatory and MUA activity in the BLAa of a pre-juvenile male mouse pup. **(ii)** Same as C (ii) for corresponding oscillatory and MUA activity in the BLAa of an adolescent male mouse pup. Note the prominent activity in the theta range (3-6, 6-12 Hz). **F, (i)** Average logarithmic power spectra of the LFP in the BLAa of male (left) and female (right) control (grey) or ELS (blue or magenta) pre-juvenile mice (males, control: N = 13, ELS: N = 14, females, control: N = 12, ELS: N = 12). Note the peaks in theta range (3-5 and 6-8 Hz). **(ii)** Same as F (i) for adolescent control and ELS male and female mice (males, control: N = 12, ELS: N = 13, females, control: N = 13, ELS: N = 17). Note the prominent broad peak in low theta activity (3-6 Hz).

#### Recording protocol

After a 20-25 min recovery period, simultaneous recordings of local field potential (LFP) and multi-unit activity (MUA) were performed from PL and IL subregions of the mPFC and the BLAa. The position of recording sites over PL, IL, and BLAa was confirmed by post-mortem histological evaluation. Both LFP and MUA were recorded at a sampling rate of 32 or 30 kHz using a multi-channel extracellular amplifier (Digital Lynx 4SX, Neuralynx, USA or Open Ephys acquisition board, Open Ephys, USA) and their corresponding acquisition software. During recording the signal was band-pass filtered between 0.1 Hz and 8 kHz (Neuralynx) or 0.1 Hz and 7.9 kHz (Open Ephys).

#### Perfusion/Fixation

After the recording, pups received another dose of more concentrated urethane (2g/kg) for deep surgical anaesthesia and were perfused with 0.1 M PBS followed by 4% paraformaldehyde (PFA). Brains were postfixed in 4% PFA for a further 48 h. Free-floating 70 µm coronal sections containing mPFC and BLAa were cut on a vibratome (Leica VT 1200S, Leica Biosystems, USA) and mounted onto adhesive slides (SuperFrost Plus, VWR, USA). Sections containing the labelled electrode tracts were imaged using brightfield microscopy (Olympus BX61, Olympus, Japan) and a 1,25x objective connected to a camera (Olympus DP73, Olympus, Japan).

#### Data analysis

Channels for analysis were selected on the basis of post-mortem histological investigation, i.e. which recording sites of fluorescently-marked electrodes were confined to PL, IL, and BLAa (n = 182 animals in total). A matching atlas plate from the adult mouse Allen Brain Reference Atlas (Allen Mouse Brain Atlas, atlas.brain-map.org) was superimposed at a similar rostral-caudal level on the brightfield photomicrograph image to determine which recording electrodes were located within the regions of interest (Figure 1B). Of those, individual channels with best signal-to-noise ratio over superficial layers II/II or deep layers V/VI of PL or IL, and medial aspects of predominantly BLAa were analysed according to the anatomical development of this projection ^28^. Recordings where the electrode placement was outside the areas of interest, or showing prominent heartbeat or noise artifacts, were excluded from further analysis. Data were imported and analysed off-line using custom-written MATLAB scripts (v. R2023a, MathWorks, USA). The signals were low-pass filtered (<1000 Hz) using a third-order Butterworth filter before downsampling to 2000 Hz. An IIR notch filter was applied at 50 Hz to attenuate line noise. All data analyses were performed blind to the group belonging of the pup.

#### Spectral analysis

The power spectral density (PSD) was estimated using Welch’s method (MATLAB function “*pwelch*”) and a sliding window approach (10-s Hamming window, 50% overlap).

#### Coherence analysis

To measure spectral correlation between signals in the mPFC and BLAa, magnitude-squared coherence (MSC) was calculated with the MATALB function *mscohere*. MSC is defined as a function of the PSD of each signal (*Pxx(f) and Pyy(f),* where x and y represent signals from different areas), and the cross power spectral density *Pxy(f)*, formally related as ***Equation 1***. Data was bandpass-filtered between 0-100 Hz and divided into 3-second windows with 50% overlap for this analysis. Confidence intervals were determined by repeating MSC calculation between a 3-second window in one area, shuffled with a Monte Carlo simulation (100 iterations per window), and the corresponding original window in the other area. The analysis was repeated for all windows initially used for the calculation of the MSC, and the 95^th^ percentile of the resulting distribution was used as confidence limit. To compare the coherence between groups, the area under the curve in the band of interest (low-theta, 3-5 Hz) was calculated for each experiment with trapezoidal numerical integration.

#### Spike sorting

All channels within PL, IL or BLAa containing spiking activity were imported into the Offline Sorter^TM^ software (Plexon) to sort MUA into single units. First, the data was high-pass filtered (>400 Hz) using a 2^nd^ order Butterworth filter. Then, spikes were detected by using amplitude threshold that was set manually after visual inspection, and the spike timestamp was defined as the largest amplitude event within each electrode. The detected spikes were automatically sorted into individual units using the Valley Seeking or K-means method. The Valley Seeking method applies an algorithm to inter-point distances to detect the number of clusters and cluster membership. The K-means method is an iterative algorithm that assigns the waveforms to one of the user-indicated cluster centres, based on Euclidean distance in feature space. Units with a signal-to-noise ratio (SNR) below 5 for OpenEphys recordings or below 11 for Neuralynx recordings were excluded. Individual spiking units were identified if their corresponding spike clusters were clearly separated in feature space (Isolation Distance > 25 and L-Ratio < 0.5), the waveform shape was consistent with a clear resemblance to derivatives of action potentials and the interspike interval (ISI) histogram showed a refractory period of > 1 ms (Plexon Offline Sorter). If a unit occurred simultaneously in more than one channel, it was included only in the channel with the highest SNR, to prevent double counting.

#### Analysis of single unit firing activity

The mean firing rate of each unit (number of spikes/s) and the Fano factor (variance of number of spikes/average of number of spikes) were calculated in MATLAB using the exported spike timestamps from the Plexon Offline Sorter.

#### Sorting of units into putative excitatory regular spiking vs putative inhibitory fast-spiking

To sort the recorded units into putative excitatory vs putative inhibitory, we used the spike waveforms and plotted the valley to peak latency as a function of the half-width (see scatter plots in Suppl. Fig. 6A), as previously used in the developing mPFC ^31, 32^. Regular spiking vs fast spiking units were distinguished by setting a manual threshold (half-width < 0.3 ms, valley-to-peak time < 0.3 ms).

#### Phaselocking analysis

To study spike-field synchronization (i.e. locking of spike time to LFP phase), we used the LFP Analysis toolbox (Beteje toolbox by Jelfs, https://github.com/beteje/LFP_Analysis,)^33^, along with custom-written MATLAB scripts (available upon request). Single-unit spike timestamps obtained with Plexon were imported into MATLAB, along with the corresponding LFP filtered between 0-100 Hz. The phase of the LFP at each spike time was estimated with a continuous wavelet transform (Morlet, 4 cycles) for 5 frequencies between 3 and 5 Hz, in 0.5 Hz intervals. Phase 0° corresponded to the peak of the oscillation, whereas phase ±180° to the through of the oscillation. To verify whether a unit was significantly phase-locked to the LFP, a Raileigh test with a threshold of 0.05 with Bonferroni correction based on the number of frequencies analysed (p = 0.05/5 = 0.01) was performed. A unit was deemed to be locked to the LFP if a significant Bonferroni-corrected p value was obtained for at least one frequency in the 3-5 Hz range. The preferred phase angle for each group was calculated as the circular mean, in degrees, of the significantly locked units, using the CircStat toolbox in MATLAB ^33^. The difference in the average phase angle between groups was measured with the Watson-William test. The difference in distribution between locked and non-locked units between groups was evaluated with the Fisher’s test.

### 2.4 Analysis of ΔFosB-positive neurons in BLA

#### Tissue preparation

Male and female control and ELS mice were sacrificed at two different time points, at P14, immediately after the LBN treatment, and at P18, corresponding to the time of the *in vivo* electrophysiological recordings (control: P14, male: N= 5, female: N= 5, P18, male: N= 5, female: N = 4, from 7 litters in total; ELS: P14, male: N = 5, female: N = 3, P18, male: N = 5, female: N = 4, from 4 litters in total). Mice were deeply anesthetized with urethane (2g/kg, Sigma-Aldrich, USA) and were intracardially perfused with 4% PFA in 0.1 M PBS for 20 minutes. Following perfusion, brains were extracted and stored in 4% PFA for 48 h at 4°C. Then, brains were washed with 0.1 M PBS and free-floating 30 µm coronal sections containing BLAa were cut on a vibratome (Leica VT 1200S, Leica Biosystems, USA). Sections were stored in anti-freeze solution (30% ethylene glycol, 30% glycerol, 30% distilled water and 10% 0.244 M phosphate buffer 2xPO4) at -20°C for further processing.

#### Immunohistochemistry

Brain sections were washed for 5 x 7 min in PBS + 0.1 % Triton X-100 (PBS-T) and incubated in blocking solution (PBS-T + 5% donkey serum) for 2 h at room temperature (RT). Next, sections were incubated in PBS-T containing the primary antibodies (see Table 1) overnight at 4°C. Then, after washing 3 x 15 min in PBS-T, the sections were incubated for 2 h at RT in PBS-T containing the secondary antibodies. Finally, sections were washed 3 x 15 min in PBS-T and mounted using Fluoromount-G^TM^ mounting medium with DAPI (Thermo Fisher).

**Table 1.** Primary and secondary antibodies used in this study.

| Immunogen | Dilution | Host | Manufacturer | ID |
| --- | --- | --- | --- | --- |
| PanFosB<br>(FosB/ΔFosB) | 1:300 | Mouse | Abcam | Ab11959 |
| FosB | 1:300 | Rabbit | Thermo Fisher | PA5-79280 |
| GAD67/GAD1 | 1:1000 | Goat | Abcam | Ab80589 |
| Mouse-488 | 1:2000 | Donkey | Abcam | Ab150105 |
| Rabbit-568 | 1:2000 | Donkey | Abcam | Ab175470 |
| Goat-647 | 1:1000 | Donkey | Abcam | Ab150131 |

#### Light microscopy imaging

Light microscopy images from four brain sections/animal containing BLAa were obtained with the Zeiss Axio Imager M2 microscope equipped with an Apotome 2 structured illumination microscopy (SIM) slider, using Plan Apo 20x (NA = 0.8) objective (Zeiss, Oberkochen, Germany), fitted with a Hamamatsu ORCA Flash 4.0 V2 camera (Hamamatsu Photonics K.K., Japan). The ZEN 2 software (Zeiss) was used for image acquisition.

#### Image analysis

To estimate the density of ΔFosB immunopositive (+) cells in the BLA, images with merged channels for FosB and PanFosB (FosB/ΔFosB) were analysed using ImageJ Fiji software. FosB^+^ cells as well as FosB/PanFosB double positive neurons were excluded and only the remaining PanFosB^+^ within the boundaries of BLAa were manually counted, as only those expressed the Δ splice variant of FosB. To quantify the intensity of the ΔFosB fluorescent signal, the difference between the single cell signal and the section’s background was determined. The quantification was then normalized to the average intensity values measured in controls ^35^. The density estimation of GAD67^+^ cells and the quantification of the GAD67 signal intensity were conducted using the same methods.

### 2.5 Statistical analysis

All statistical analyses were done on raw data using Prism, version 9.4.0 (GraphPad, USA), custom-written MATLAB scripts, or MATLAB community toolboxes. The sample size was based on previous experience on similar experiments. The distribution of the data was tested with the D’Agostino-Pearson test, and two-sided parametric (unpaired t-test, Welch’s t-test) or non-parametric tests (Mann-Whitney U test) were chosen accordingly. Uniformity of circular data was measured with the Rayleigh test. Differences between groups in circular data were evaluated with the Watson-Williams test. To measure the relative effect of the LBN treatment onto any outcome along the two time points (P18-20 and P43-47), a two-way ANOVA was used, with post-hoc Sidák correction for multiple comparisons. The results were considered significant when *p* < 0.05. All the pooled data are given as mean ± SEM or or mean ± SD or median (interquartile range (IQR)) as noted. In the figures, statistical significance levels are indicated by asterisks as follows: *p < 0.05, **p < 0.01, ***p < 0.001, and ****p < 0.0001.

## 3. Results

### 3.1 ELS does not affect the development of local oscillatory network activity in mPFC or BLAa

To test the effect of ELS on the development of prefrontal-amygdala networks, male and female pups were exposed to the ELS LBN paradigm between P4-14 with additional MS for 1 h at P8, P10, and P12 (Fig. 1A). This combined treatment of LBN+MS has been shown to result in an amygdala phenotype before ^15^. Previous studies characterizing the LBN model have shown a consistent decrease in physical growth i.e. body weight or body weight gain of LBN pups compared to controls^12, 14, 36^. Similarly, we also found a significant decrease in body weight in ELS males of 19.87 ± 1.69 % at the end of the LBN treatment period (P14, Suppl. Fig. 1Ai), which slowly recovered to 12.84 ± 1.62 % during pre-juvenile (P18-21) development and to 6.48 ± 2.40 % during adolescence (P43-47, Two-Way ANOVA, factor treatment: F(2, 179) = 35.00, p < 0.0001, factor age: F(2, 179) = 1936, p < 0.0001, n = 18-46 pups/group, Sidak’s multiple comparisons tests: P14, p < 0.0001; P18, p = 0.025; P45, p = 0.0025). ELS females also showed a significant decrease in body weight at P14 (13.79 ± 2.30 %, Suppl. Fig. 1Aii), as well as during pre-juvenile (10.88 ± 1.73 %) and adolescent development (6.44 ± 1.42 %, Two-Way ANOVA, factor treatment: F(2, 174) = 30.19, p < 0.0001, factor age: F(2, 174) = 1871, p < 0.0001, n = 24-45 pups/group, Sidak’s multiple comparisons tests: P14, p = 0.024; P18, p = 0.0085; P45, p = 0.0005). Hence, we are confident that our implementation of the LBN ELS paradigm was successful.

Following the ELS paradigm, spontaneous local-field potential (LFP) and multi-unit activity (MUA) within superficial (II/III) or deep (V/VI) layers of the right PL or IL cortex (Fig. 1Bi, Suppl. Fig. 1Bi) or the BLAa (Fig. 1Bii, Suppl. Fig. 1Bii) were recorded from pre-juvenile (P18-20) or adolescent (P43-47) male and female ELS pups under light urethane anaesthesia as well as from control pups at the same age.

Spontaneous prefrontal network activity in control male and female pre-juvenile mice (Fig. 1Ci, Di, Supp. Fig. 1Ci) was continuous and was characterized by prominent high-amplitude delta rhythms (0.3-0.5 Hz) superimposed by theta, most prominent low-theta (3-6 Hz), beta and gamma (12-100 Hz) activity as well as local MUA, as typical for urethane-induced slow-wave sleep-like activity ^37–39^. During adolescent development (Fig. 1Cii), high-amplitude delta activity became faster (peak frequency ∼1 Hz) and particularly the superimposed low-theta activity (3-6 Hz) was even more prominent, evident as clear peaks in the related power spectra (Fig. 1Dii). Network activity was similar between PL and IL (Fig. 1D, Suppl. Fig. 1C) and between superficial and deep cortical layers within either developmental period. No differences were found between males and females at either age. Furthermore, cortical network activity in ELS mice was also not significantly different from controls in neither males nor females at either developmental time point (Fig 1D).

Spontaneous network activity in pre-juvenile BLAa was also characterized by high-amplitude delta activity (0.3-0.5 Hz) with superimposed prominent theta activity, particularly low-theta (3-6 Hz), as well as beta and gamma activity and MUA (Fig. 1Ei, Fi). Similarly to cortical network activity, during adolescent development delta rhythms became faster (∼ 1 Hz, Fig. 1Eii) and low-theta activity increased in power, as it can be seen by the prominent peaks (3-6 Hz) in the power spectrum (Fig. 1Fii). Network activity in the BLAa was similar between males and females at either developmental time point. Moreover, BLAa network activity was also not significantly different between ELS and control mice neither during pre-juvenile nor during adolescent development (Fig. 1F).

Hence, it seems that ELS does not affect the generation of neither cortical nor amygdala spontaneous network activity per se during neither pre-juvenile nor adolescent development.

### 3.2. ELS impairs prefrontal-amygdala oscillatory coupling during pre-juvenile development in males only

ELS has been shown to impair functional interactions within developing prefrontal-amygdala networks ^15, 21–26^. However, most of the data was obtained by resting-state fMRI and electrophysiological investigations in mouse models of ELS remain sparse. Our previous study in pre-pre-juvenile (P14-15) ELS rats exposed to daily maternal separation between P2-14 revealed a decrease in oscillatory coupling within prefrontal-amygdala networks in the low-theta range (2-5 Hz) in males only ^26^. To investigate if sex-dependent impairments in functional coupling within prefrontal-amygdala networks particularly in low-theta range is a general signature of ELS across different species and ELS models, we analysed coherence as a measure of coupling synchrony in pre-juvenile and adolescent control and ELS male and female mice. Generally, coherence in the 0-40 Hz range was significantly larger compared to shuffled data within both prelimbic superficial layers-BLAa or prelimbic deep layers-BLAa in pre-juvenile males and females (Fig. 2A), and the same was true for coherence within pre-juvenile infralimbic-BLAa networks (Suppl. Fig. 2). Since our previous study in developing rats identified low-theta as a substrate for prefrontal-amygdala interactions ^26^, we focussed our coherence analysis on the low-theta range (3-5 Hz) as well. Coherence within either prelimbic superficial layers-BLAa or prelimbic deep layers-BLAa networks showed a broad peak in low-theta range (3-5 Hz) in both pre-juvenile control male and female pups (Fig. 2A). In ELS mice, this peak in low-theta range was even more prominent in both pre-juvenile males and females. However, when comparing the area under the curve (AUC) from 3-5 Hz between treatment groups, low-theta coherence was selectively increased versus controls only within prelimbic superficial layers-BLAa networks and only in pre-juvenile males (Fig. 2B, t(21) = 2.297, p = 0.032, N = 11-12, unpaired t-test). In contrast, no significant changes in low-theta coherence were observed in pre-juvenile ELS females. Furthermore, when analysing low-theta coherence within infralimbic-BLAa networks (Suppl. Fig. 2), no significant changes were observed neither within infralimbic superficial layers-BLAa nor within infralimbic deep layers-BLAa networks, in neither pre-juvenile ELS males nor females versus controls.

**Figure 2.**
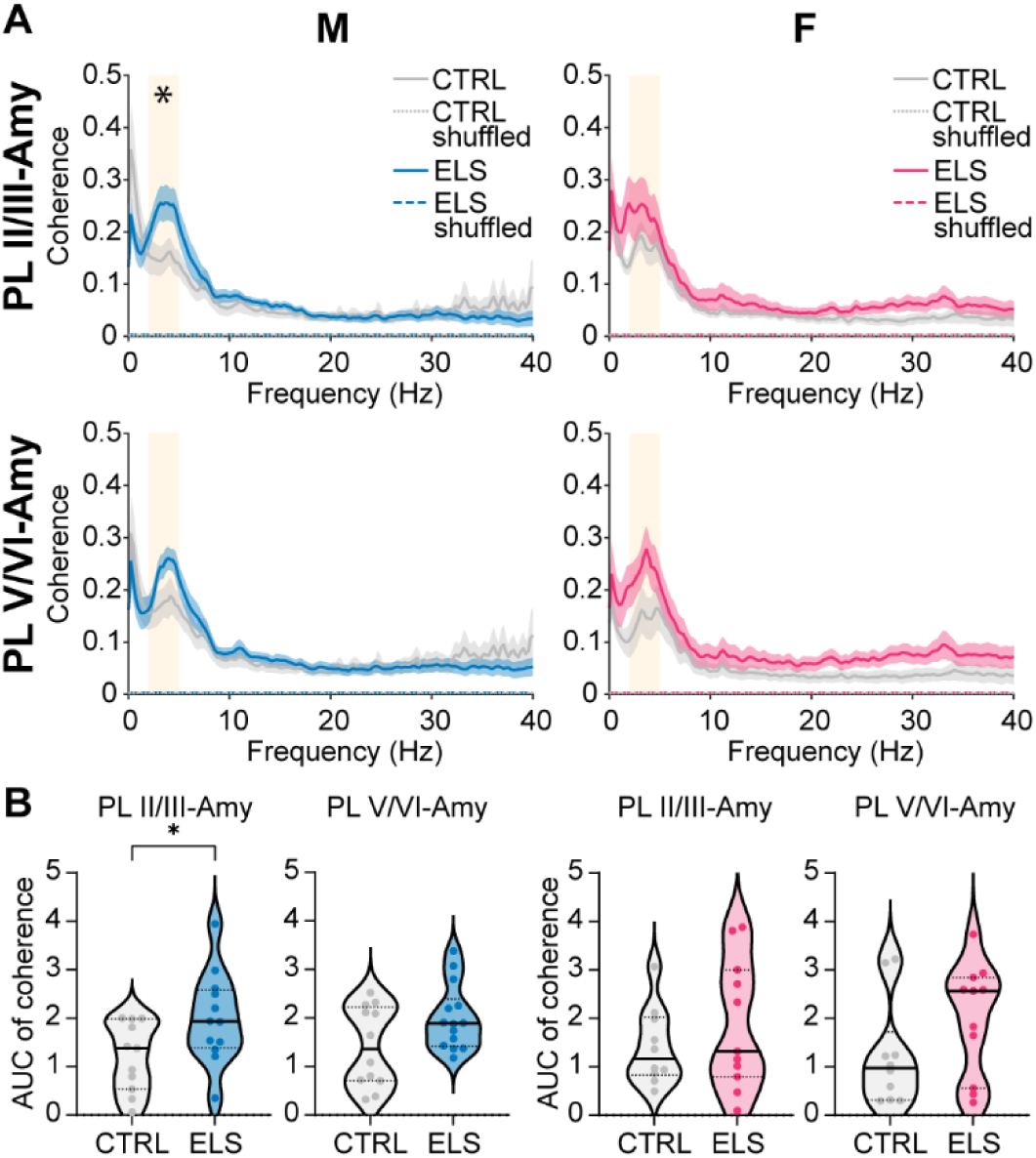
ELS leads to hypercoupling within prelimbic-amygdala networks in male pre-juvenile mice. A, Average coherence spectra of LFP in PL, layers II/III and LFP in BLAa (top) or LFP in PL, layers V/VI and LFP in BLAa (bottom) of male (left) and female (right) control (grey) or ELS (blue or magenta) pre-juvenile mice (males: N = 11-14/group, females: N = 10-11/group). Note the broad peak in coherence in the low-theta range (3-5 Hz) in all groups, which is particularly increased for coherence within PL layers II/III - BLAa networks in male ELS mice. **B,** Violin plots with median and IQR showing the area under the curve (AUC) for coherence in the low-theta band (3-5 Hz) within prelimbic layer II/III - BLAa or prelimbic layer V/VI - BLAa networks in male (left) and female (right) control (grey) or ELS (blue or magenta) pre-juvenile mice. Note the significant increase in low theta coherence particularly within prelimbic layer II/III - BLAa networks in male pre-juvenile ELS mice. See Suppl. Table 3 for the exact number of pups per group in each graph. *p < 0.05, unpaired t-test.

Hence, ELS selectively impairs long-range functional interactions within pre-juvenile prelimbic superficial layers-BLAa networks in a sex-dependent manner only in males while females were not significantly affected.

### 3.3. Prefrontal-amygdala oscillatory coupling is not affected during adolescent development

To assess if the impairments in functional long-range interactions within prelimbic-amygdala networks persist into adolescence, we computed coherence within prelimbic-BLAa or infralimbic-BLAa networks in adolescent male and female mice. Again, we focussed our analysis on oscillatory coupling within low-theta (3-5 Hz) range. Control adolescent male and female mice showed prominent peaks in low-theta coherence within prelimbic superficial layers-BLAa or prelimbic deep layers-BLAa (Fig. 3A, B). However, to our surprise the pre-juvenile hypercoupling within prelimbic superficial layers-BLAa was not present anymore in adolescent ELS males, and low-theta coherence within this network was similar to control males. Likewise, ELS did not affect low-theta coherence within prelimbic superficial layers-BLAa in adolescent females. Also, low-theta coherence within prelimbic deep layers-BLAa was similar between control and ELS mice in both males and females.

**Figure 3.**
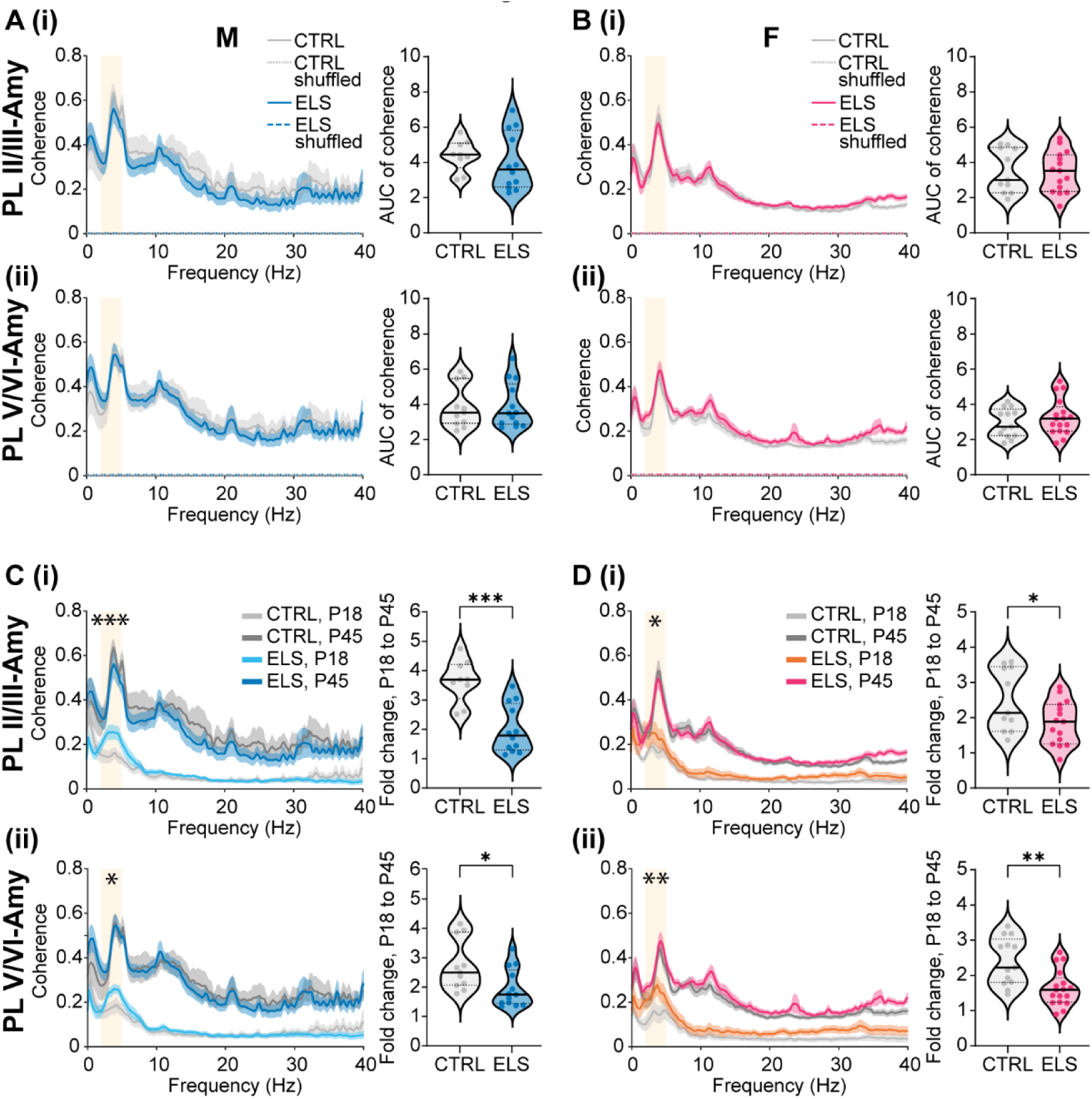
ELS does not affect oscillatory coupling within adolescent prefrontal-amygdala networks. A, **(i)** Average coherence spectra of LFP in PL, layers II/III and LFP in BLAa, and corresponding violin plots with median and IQR showing the area under the curve (AUC) for coherence in the low-theta band (3-5 Hz) in adolescent control (grey, N = 9) and ELS (blue, N = 12) male mice. **(ii)** Same as A (i) for coherence between the LFP in PL, layers V/VI and BLAa (control, N = 11, ELS, N = 13). **B, (i)** Same as A (i) for adolescent control (grey, N = 11) and ELS (magenta, N = 15) female mice. **(ii)** Same as A (ii) for adolescent control (grey, N = 13) and ELS (magenta, N=17) female mice. **C, (i)**, Average coherence spectra between the LFP in PL, layers II/III and the LFP in BLAa for control pre-juvenile (light grey) and adolescent (dark grey) as well as ELS pre-juvenile (cyan) and adolescent (blue) male mice. The corresponding violin plots with median and IQR (right) show the fold change in low-theta coherence (3-5 Hz) between pre-juvenile and adolescent control (grey, N = 9) or pre-juvenile and adolescent ELS (blue, N = 12) male mice. Note that the fold increase in low-theta coherence is significantly smaller in ELS mice, suggesting that ELS promotes precocious development of functional interactions within these networks. **(ii)** Same as C (i) for coherence between the LFP in PL, layers V/VI and the LFP in BLAa (control: N = 11, ELS: N = 13). **D, (i)** Same as C (i) for female pre-juvenile (light grey) and adolescent (dark grey) control and pre-juvenile (orange) and adolescent (magenta) ELS mice. The corresponding violin plots with median and IQR (right) show the fold change in low-theta coherence (3-5 Hz) between pre-juvenile and adolescent control (grey, N = 11) or pre-juvenile and adolescent ELS (magenta, N = 15) female mice. **(ii)** Same as D (i) for coherence between the LFP in PL, layers V/VI and the LFP in BLAa (control: N = 13, ELS: N = 17). *p < 0.05, **p < 0.01, ***p < 0.001, unpaired t-test.

Coherence within infralimbic superficial layers-BLAa or infralimbic deep layers-BLAa also showed prominent peaks within low-theta range (Suppl. Fig. 3A, B) in both male and female adolescent control mice. ELS did not affect low-theta coherence within infralimbic-BLAa networks either. Low-theta coherence in adolescent ELS males or females within infralimbic superficial layers-BLAa or infralimbic deep layers-BLAa was not significantly different from controls.

Hence, during adolescence, spontaneous oscillatory coupling within prelimbic-BLAa or infralimbic BLAa networks was not affected by ELS in neither male nor female mice.

### 3.4 ELS mice show a smaller developmental increase in mPFC-amygdala coherence indicating a precocious development in coupling strength

It has been shown both in humans with a history of ELS and in rodent models that ELS causes a precocious development of amygdala networks as well as prefrontal-amygdala structural and functional connectivity ^6, 25, 40^. Here we analysed if ELS also has an effect on the developmental increase in oscillatory coupling within prefrontal-amygdala networks. Generally, coherence across 0-40 Hz Hz within both prelimbic-BLAa (Fig. 3C, D) or infralimbic-BLAa (Suppl. Fig. 3C, D) networks was higher in adolescent versus pre-juvenile mice. When calculating the fold-change in low-theta coherence between pre-juvenile and adolescent mice, low-theta coherence increased by 3.7 (1.2) fold within prelimbic superficial layers-BLAa or 2.5 (1.8) fold within prelimbic deep layers-BLAa in control males. This change was significantly smaller in ELS mice, with only a 1.8 (1.6) fold increase in coherence within prelimbic superficial layers-BLAa (t(19) = 4.76, p = 0.0001, N = 9-12, unpaired t-test) or a 1.7 (1.2) fold increase in coherence within prelimbic deep layers-BLAa (t(22) = 2.55, p = 0.018, N = 11-13, unpaired t-test). In control females, low-theta coherence increased by 2.1 (1.8) fold within prelimbic superficial layers-BLAa or by 2.2 (1.2) fold within prelimbic deep layers-BLAa. Similar to ELS males, the increase in low-theta coherence was significantly smaller in ELS females and increased only by 1.9 (1.1) fold within prelimbic superficial layers-BLAa (t(24) = 2.25, p = 0.034, N = 11-15, unpaired t-test) or by 1.6 (0.7) fold within prelimbic deep layers-BLAa (t(28) = 3.55, p = 0.0014, N = 13-17, unpaired t-test). Also, infralimbic-amygdala low-theta coherence was higher in adolescent vs pre-juvenile mice across the 0-40 Hz spectrum (Suppl. Fig. 3C, D). Low-theta coherence increased by 3.1 (0.5) fold within infralimbic superficial layers-BLAa or by 2.8 (1.2) fold within infralimbic deep layers-BLAa in control males. Interestingly, ELS only led to a significant decrease in this change, suggesting a precocious development within infralimbic superficial layers-BLAa (ELS: 2.1 (1.2), U = 19, p = 0.035, N = 9-10, Mann-Whitney test) but not within infralimbic deep layers-BLAa in male mice. In control females, low-theta coherence increased by 2.2 (1.3) fold within infralimbic superficial layers-BLAa or 2.3 (1.7) fold within infralimbic deep layers-BLAa. Remarkably, the increase in theta coupling within infralimbic-amygdala networks across development was not affected by ELS.

Accordingly, the largest difference between control and ELS mice regarding the increase in low-theta coupling was within prelimbic superficial layers-BLAa in males (Fig, 3Ci) due to the particularly strong increase in theta coupling within controls across development, as well as the prominent low-theta hypercoupling within this network in pre-juvenile ELS males. Hence, the increase in low-theta coherence across development was particularly diminished within prelimbic-amygdala networks in both ELS males and females. This may indicate that ELS causes a precocious development in functional long-range interactions predominantly within prelimbic-amygdala networks in this ELS mouse model.

### 3.5 ELS leads to a higher firing activity of a subpopulation of units in pre-juvenile BLAa and to a widespread increase in firing activity in adolescent BLAa

One of the hallmark signatures of ELS in humans is an increased amygdala reactivity ^4, 41^. Previous work in mouse models of ELS also suggested an increased activity of BLA neurons based on cFos analysis, ^17^. Here, we analysed the firing activity in BLAa in male and female control and ELS mice. For this, MUA was sorted into single units using the Plexon Offline Sorter. In pre-juvenile control males, n = 41 BLAa single units from N = 12 mice were obtained. Their firing rates were low 0.62 (1.34) Hz and the Fano factor, a measure for the variability of neural spike trains, was 1.86 (1.00) suggesting that spiking patterns were irregular and variable over time (Fig. 4A). In pre-juvenile ELS males, n = 36 single BLAa units from N = 12 mice were obtained. Their firing rates (0.68 (1.95)) and Fano factors (1.83 (1.03)) were similar to controls (Fig. 4A). However, when plotting the logarithmic frequency distribution of firing rates (Fig. 4B), control mice showed a unimodal distribution while ELS mice showed a bimodal distribution of firing rates. This suggests that in controls the data was obtained from a single, similarly active population of neurons. Interestingly, the bimodal distribution in ELS mice indicates two distinct populations of neurons, one firing at higher rates than the other. We did not find any anti-correlation between firing rate and half-width (Suppl. Fig. 4Ai) which would have been expected if the more active population of neurons after ELS were fast-spiking interneurons. Clustering using similar criteria as for the mPFC (see below) did not reveal more than one cluster per group, which were also overlapping (Suppl. Fig. 4Aii). Hence, this suggests that increased neuronal activity in the BLAa after ELS is due to activation of a subpopulation of principal neurons rather than increased population activity of fast-spiking interneurons. Finally, we also analysed and compared the spike waveforms in both groups. The valley-to-peak time (ms) of the spikes, corresponding to the duration of the medium afterhyperpolarization (mAHP) of the extracellular action potential, was significantly decreased in ELS males versus control males (Fig. 4Ci, control: 0.40 (0.1), ELS: 0.32 (0.1), t(75) = 4.262, p < 0.0001, n = 36-41, unpaired t-test). This suggests a significantly shorter mAHP in the spikes of ELS mice, which is a typical indicator of increased excitability ^42^. When computing the full width at half maximum (FWHM) of the waveform (see diagram in Fig. 4Ci), the duration in control and ELS mice was similar (Fig. 4Cii), suggesting, in line with our results from clustering, recordings were sampled from similar types of neurons in both groups, most likely predominantly BLAa principal neurons.

**Figure 4.**
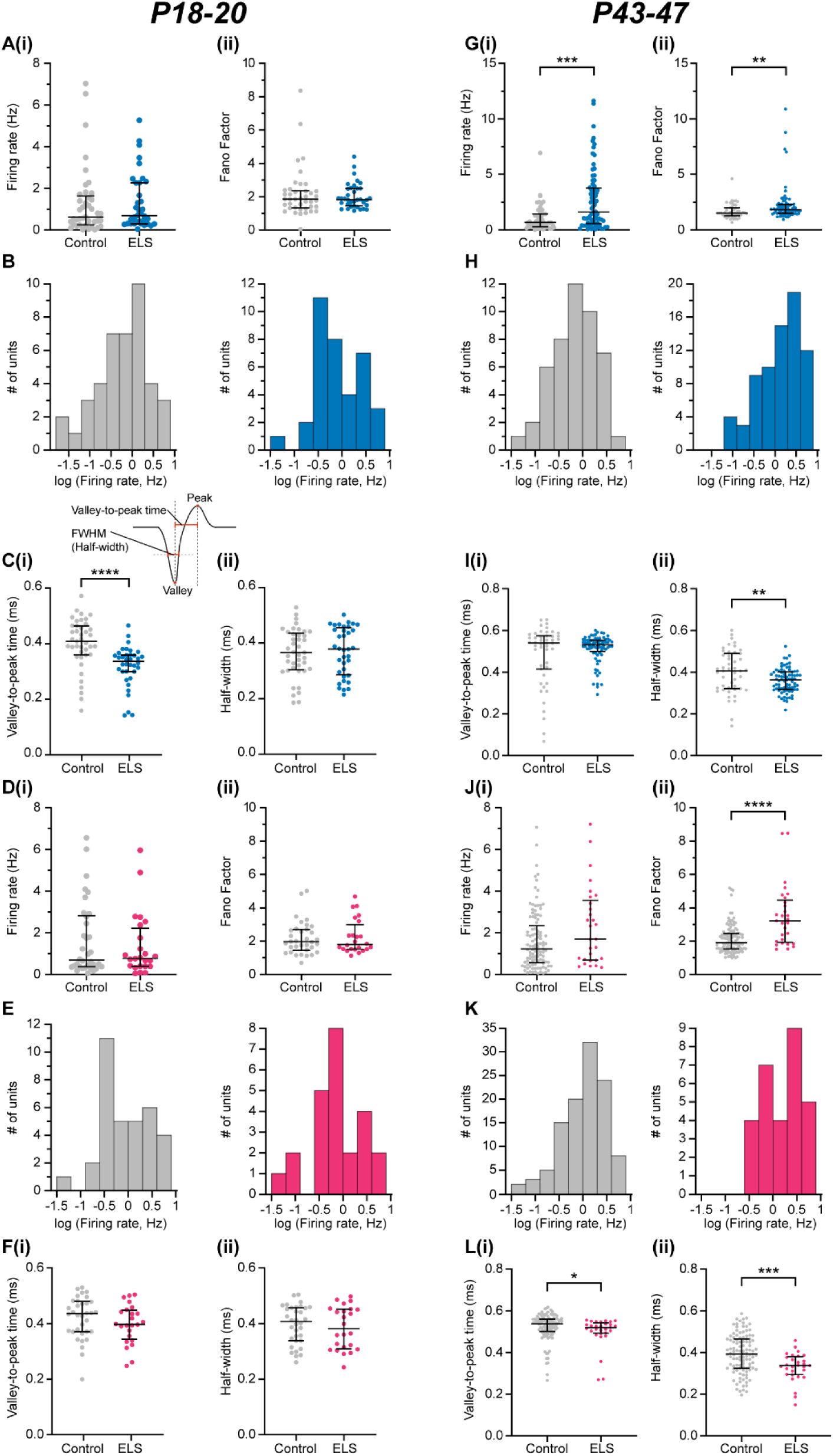
ELS increases the firing rates of a subpopulation of neurons in the BLAa in both male and female pre-juvenile mice and leads to a widespread increase in mean firing rates in BLAa of adolescent males. **A,** **(i)** Distribution and median (IQR) of instant firing rates (Hz) of single units in the BLAa in control (grey, n = 41 units from N = 12 pups) and ELS (blue, n = 36 units from N = 12 pups) male pre-juvenile mice. **(ii)**, Same as A (i) for Fano factor. **B**, Logarithmic frequency distribution (bin width 0.3 s) of firing rates of single units in BLAa from both control (grey, n = 41 units, left) and ELS (blue, n = 36 units, right) pre-juvenile male mice. Note the bimodal distribution of firing rates in ELS mice, suggesting increased firing rates in a subpopulation of BLA neurons. **C, (i)** Distribution and mean ± SD of the mean valley-to-peak time (ms) corresponding to the mAHP of the extracellular action potential (see diagram above) of single units in BLAa in control (grey, n = 41 units) and ELS (blue, n=36 units) male pre-juvenile mice. Note the prominent decrease in ELS male mice. The diagram shows a representative action potential with key parameters highlighted in red. **(ii)** Same as C (i) for the full width at half maximum (FWHM, half-width (ms), see diagram in C (i)) of the extracellular action potential waveform. **D, (i)** Distribution and median (IQR) of instant firing rates (Hz) of single units in the BLAa in control (grey, n = 34 units from N = 9 pups) and ELS (magenta, n = 24 units from N = 10 pups) female pre-juvenile mice. **(ii)**, Same as C (i) for Fano factor. **E,** Logarithmic frequency distribution (bin width 0.3 s) of firing rates of single units in BLAa from both control (grey, n = 34 units, left) and ELS (magenta, n = 24 units, right) pre-juvenile female mice. Note the bimodal distribution of firing rates in ELS mice, suggesting increased firing rates in a subpopulation of BLA neurons. **F, (i)** Distribution and mean ± SD of the mean valley-to-peak time (ms) of single units in BLAa in control (grey, n = 36 units) and ELS (magenta, n = 24 units) female pre-juvenile mice. **(ii)** Same as F (i) for the full width at half maximum (ms) of the extracellular action potential waveform. ****p < 0.0001, Mann-Whitney test. **G, (i)** Distribution and median (IQR) of instant firing rates (Hz) of single units in the BLAa in control (grey, n = 47 units from N = 10 pups) and ELS (blue, n = 77 units from N = 11 pups) male adolescent mice. Mann-Whitney test, ***p = 0.0004. **(ii)**, Same as G (i) for Fano factor. Mann-Whitney test, **p = 0.0026. **H**, Logarithmic frequency distribution (bin width 0.3 s) of firing rates of single units in BLAa from both control (grey, n = 47 units, left) and ELS (blue, n = 77 units, right) adolescent male mice. **I, (i)** Distribution and median (IQR) of the mean valley-to-peak time (ms) corresponding to the mAHP of the extracellular action potential of single units in BLAa in control (grey, n = 47 units) and ELS (blue, n = 77 units) male adolescent mice. Unpaired t-test, **p = 0.0041. **(ii)** Distribution and mean ± SD for the full width at half maximum (FWHM, half-width (ms), see diagram in C (i)) of the extracellular action potential waveform. **J, (i)** Distribution and median (IQR) of instant firing rates (Hz) of single units in the BLAa in control (grey, n = 109 units from N = 14 pups) and ELS (magenta, n = 29 units from N = 8 pups) female adolescent mice. **(ii)**, Same as J (i) for Fano factor. Mann-Whitney test, ****p < 0.0001. **K,** Logarithmic frequency distribution (bin width 0.3 s) of firing rates of single units in BLAa from both control (grey, n = 109 units, left) and ELS (magenta, n = 29 units, right) adolescent female mice. **L, (i)** Distribution and median (IQR) of the mean valley-to-peak time (ms) of single units in BLAa in control (grey, n = 109 units) and ELS (magenta, n = 29 units) female adolescent mice. Mann-Whitney test, *p = 0.0225 **(ii)** Distribution and mean ± SD of the full width at half maximum (ms) of the extracellular action potential waveform. Unpaired t test, ***p = 0.0006.

In pre-juvenile females, after spike sorting n = 34 BLAa single units from N = 9 control mice and n = 24 BLAa single units from N = 10 ELS mice were obtained. As in control males, baseline firing rates in the BLAa in control females (Fig. 4D) were also low (0.69 (2.41) Hz) and had a similar Fano factor (1.96 (1.22)). Furthermore, neither baseline firing rates (0.78 (1.67) Hz) nor Fano factor (1.78 (1.26)) in ELS females were significantly different from controls. However, when plotting the logarithmic frequency distribution of firing rates in both groups (Fig. 4E), similar to males, control females showed a unimodal distribution while ELS females showed a bimodal distribution of firing rates. This suggests that also in females, the data was obtained from a single similarly active population of neurons in controls, but from two separate populations in ELS mice, one with higher firing activity. Again, when clustering the spikes as for males above (Suppl. Fig. 4B), we did not find any evidence that more active neurons would be putative fast-spiking interneurons. Hence, the more active population of BLA neurons after ELS are most likely also BLA principal neurons. When computing the valley-to-peak time (ms) or the FWHM of BLAa spike waveforms (Fig. 4F), no differences were observed between control and ELS females. This suggests that in contrast to pre-juvenile ELS males, in ELS females the intrinsic excitability of BLAa neurons was not altered by ELS, but, as in males, active neurons in both groups seem to belong to similar types of cells, i.e. most likely BLAa principal neurons.

Next, we performed spike sorting and analysed neuronal firing activity in BLAa of adolescent control and ELS mice. We obtained n = 47 single units from N = 10 adolescent control males and n = 77 single units from N = 11 adolescent ELS males. Remarkably, adolescent ELS males showed a significant increase in mean baseline firing rates compared to controls (Fig. 4G(i), control: 0.71 (1.2) Hz, ELS: 1.64 (3.2) Hz, U = 1130, p = 0.0004, Mann-Whiteny test). Moreover, the Fano factor was also significantly increased in adolescent ELS males compared to controls. (Fig. 4G(ii), control: 1.53 (0.7), ELS: 1.84 (0.7), U = 1229, p = 0.0028, Mann-Whitney test). These data show that ELS not only leads to a prominent increase in baseline firing activity in males but also affects the firing patterns. The logarithmic frequency distribution of firing rates showed a unimodal distribution for both adolescent male control and ELS mice (Fig. 4H). while ELS mice exhibited a skewness towards higher firing rates in accordance with the increased firing rate (Fig. 4H). The change from a bi-modal distribution in prejuvenile ELS males to a unimodal distribution of firing rates in adolescent ELS males suggests a more widespread increase in firing activity affecting a larger population of BLAa neurons in adolescent ELS males. Similar to pre-juvenile development, we did not find any anti-correlation between firing rate and half-width (Suppl. Fig. 4Ci) and clustering also only revealed one larger cluster in each treatment group which were overlapping (Suppl. Fig. 4Cii). This suggests that increased neuronal activity in BLAa of adolescent ELS males is due to activation of predominantly on large population of neurons, proabably BLAa principal neurons. Interestingly, valley-to-peak time of the spike was not significantly different between the groups (Fig. 4I(i)), while FWHM was decreased in adolescent ELS males compared to controls (Fig. 4I(ii), t(122) = 2.93, p = 0.0041, N = 47-77, unpaired t-test).

In BLAa of adolescent females, we obtained n = 109 BLAa single units from N = 14 control and n = 29 single units from N = 8 ELS mice. In contrast to males at the same age, firing rates were not significantly increased after ELS (Fig. 4 J(i)), but the Fano factor was strongly increased (Fig. 4J(ii), control: 1.91 (0.9), ELS: 3.22 (2.5), U = 746, p < 0.0001, Mann-Whitney test). However, there was a trend towards increased firing rates (control: 1.22 (1.8), ELS: 1.70 (2.8), U = 1257, p = 0.0913, Mann-Whitney test) as well in adolescent ELS females that probably did not reach significance due to the lower number of units in this group. The logarithmic frequency distribution of firing rates showed a unimodal distribution for adolescent female controls (Fig. 4K) while the distribution remained inconclusive in adolescent ELS females due to the low number of units in this group. Both valley-to-peak time of the spike (Fig. 4 L(i), control: 0.54 (0.1), ELS: 0.52 (0.05), U = 1145, p = 0.0225) and FWHM (Fig. 4L(ii), t(136) = 3.50, p = 0.0006, unpaired t-test) were significantly decreased in adolescent ELS females compared to controls, pointing to an overall shorter duration of the action potential in this group.

Hence, while during pre-juvenile development neither the mean firing rates nor the firing patterns seem to be affected in BLAa by ELS, in both pre-juvenile ELS males and females a subpopulation of BLA neurons showed higher firing activity. Furthermore, only in pre-juvenile ELS males this was accompanied by changes in the spike waveform, suggesting an increased intrinsic excitability. During adolescent development, this phenotype was even stronger showing a prominent increase in mean firing rates in BLAa in ELS males while this effect did not reach significance in females probably due to the lower number of units. Moreover, also the firing patterns were impaired in both sexes after ELS whereas modifications in action potential shape point to altered membrane excitability or ion channel expression after ELS.

### 3.6 ELS leads to persistent activation of pre-juvenile BLA neurons in both males and females

To confirm the findings from our *in vivo* recordings, we carried out immunohistochemistry against ΔFosB, a marker of chronic activation in neurons ^43^, in the BLAa (Fig. 5). Since no antibodies against ΔFosB were available, we performed dual-immunolabelling against PanFosB, which labels both FosB and ΔFosB immunopositive (+) neurons and FosB. Those PanFosB^+^ neurons that did not colocalize with FosB were counted as ΔFosB^+^ (see Fig. 5A). Data were collected at P14, immediately after the LBN stress period, and during pre-juvenile development at P18, at the same age when the *in vivo* electrophysiological recordings were carried out. In males, two-way ANOVA revealed a significant effect of ELS treatment (F(1, 16) = 4.757, p = 0.044, N = 5 mice/group) but no effect of age nor of the age x ELS treatment interaction on the number of BLAa ΔFosB^+^ cells (Fig. 5Bi, Suppl. table 4) suggesting that ELS leads to an increase in the number of ΔFosB^+^ neurons in the BLAa in males that is independent of age. When analysing the data separately for either right or left hemisphere in both groups at both ages (Fig. 5Bii), there was still a significant effect of ELS treatment (three-way ANOVA, F(1, 29) = 6.718, p = 0.0148) on the number of ΔFosB^+^ neurons, but no effect of neither hemisphere nor age, nor any significant interactions (hemisphere x age, hemisphere x ELS treatment, age x ELS treatment, hemisphere x age x ELS treatment in males, (Suppl. table 4). We also determined the intensity of the ΔFosB signal as a percentage of control, but did not find any significant effects of neither ELS treatment nor age nor ELS treatment x age interaction (Fig. 5Biii, Suppl. table 4).

**Figure 5.**
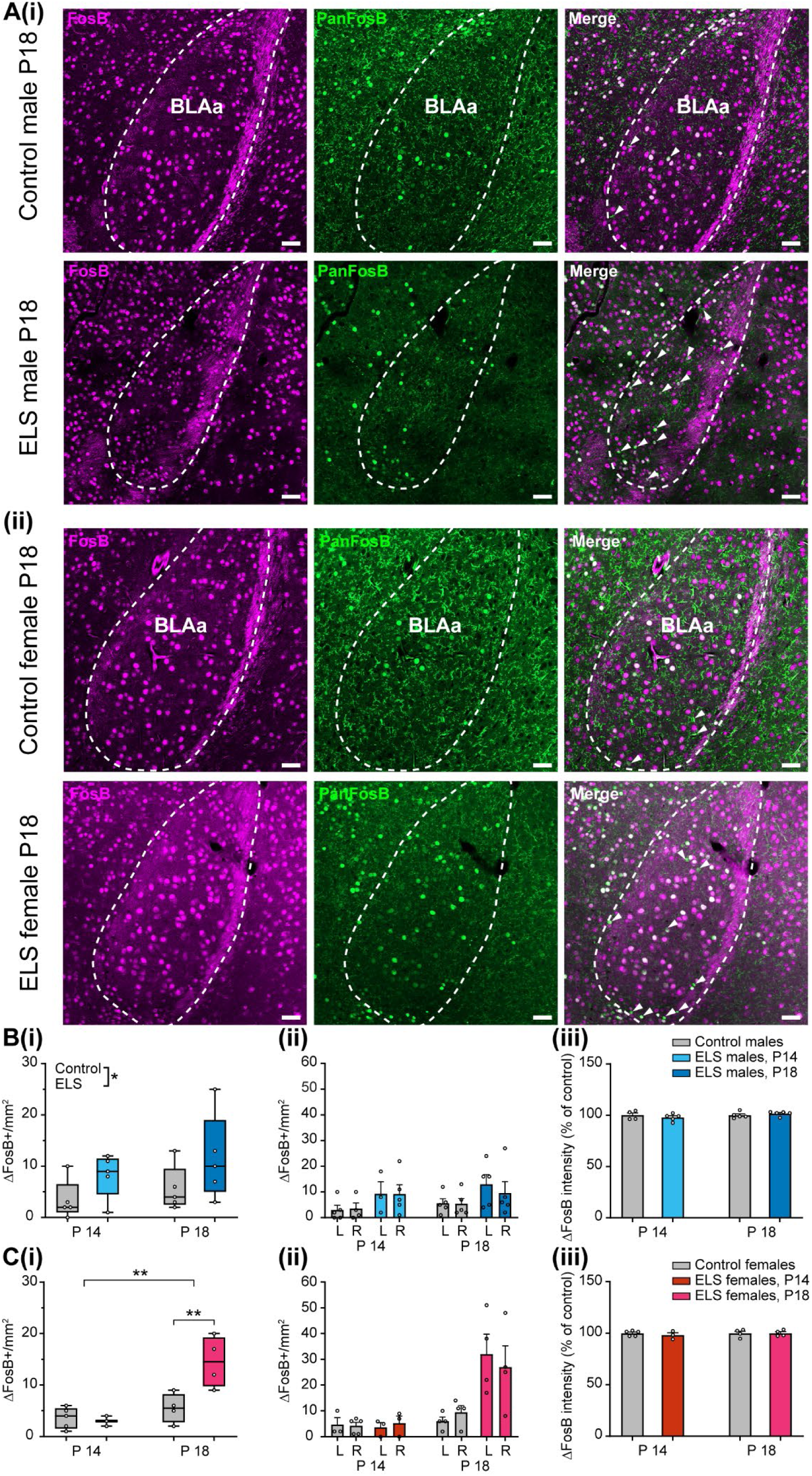
ELS causes chronic activation of BLAa neurons in both pre-juvenile males and females. A, **(i)** Representative photomicrograph images showing neurons immunopositive (+) for FosB (magenta, left) and PanFosB (green, center) in the BLAa of a control (top row) and ELS (bottom row) male P18 mouse. The merged images (right) show colocalization of FosB and PanFosB (white cells). Those PanFosB^+^ cells that do not colocalize with FosB were identified as ΔFosB^+^ (see arrowheads). **(ii)** Same as A (i) for a control and ELS female mouse. Scale bars: 50 µm. **B, (i)** Box and whisker plots of the number of ΔFosB^+^ cells/mm^2^ in the BLAa in control (grey) and ELS (blue) male mice at P14 and P18 (N = 5 mice/per group). Note the significant increase in ΔFosB^+^ cells in male ELS mice at both time points. **(ii)** Bar diagrams showing the mean ± SEM of the number of ΔFosB^+^ cells/mm^2^ in either left (L) or right (R) hemisphere in the BLAa at P14 and P18 in control (grey) and ELS (cyan or blue) male mice (N = 3-5/group). Note that there is no significant difference between the hemispheres at either time point. **(iii)** Bar diagrams showing the mean ± SEM of the ΔFosB intensity of positive neurons in the BLAa at P14 and P18 in control (grey, N = 4-5/group) and ELS (cyan or blue, N = 5/group) male mice. Note that the ΔFosB intensity is similar in both treatment groups at both time points. **C, (i)** Same as B (i) for control (grey, N = 4-5/group) and ELS (red or magenta, N = 3-4/group) female mice. Note here that the effect of ELS is age-dependent, with no change by ELS in the number of ΔFosB^+^ cells/mm^2^ at P14 but a strong significant increase by ELS at P18 in females. **(ii)** Same as B (ii) for control (grey, N = 3-5) and ELS (red or magenta, N = 3-4) female mice. Note that there is no significant difference between the hemispheres at either time point. **(iii)** Same as B (iii) in control (grey, N = 4-5/group) and ELS (red or magenta, N = 3-4/group) female mice. Note that the ΔFosB intensity is similar in both treatment groups at both time points. *p < 0.05, **p < 0.01, Two-way ANOVA with Sidak’s post-hoc test.

Interestingly, the effects of ELS in females were even stronger (Fig. 5Ci). There was a significant effect of ELS treatment on the number of BLAa ΔFosB^+^ neurons (Two-way ANOVA, F(1, 12) = 6.992, p = 0.0214, N = 3-5 mice/group, Suppl. table 4) and also a significant effect of age (F(1, 12 = 17.79, p = 0.0012) as well as an ELS treatment x age interaction (F(1, 12) = 9.132, p = 0.0106). In addition, post-hoc tests revealed a significant increase in the number of ΔFosB^+^ neurons induced by ELS treatment at P18 (p = 0.0031) but not at P14. Hence, in females ELS leads to an age-dependent increase in the number of ΔFosB+ neurons in the BLAa during pre-juvenile development at P18-20.

When analysing the data separately for left and right hemispheres (Fig. 5Cii), similar to males, there was no effect of the hemisphere, but still an effect of ELS treatment (Three-way ANOVA, F(1, 22) = 10.67, p = 0.0035, Suppl. table 4) and age (F(1, 22) = 17.98, p = 0.0003) as well as ELS treatment x age interaction (F(1, 22) = 10.54, p = 0.0037). Finally, when determining the intensity of the ΔFosB signal as a percentage of control (Fig. 5Ciii), there was no effect of ELS treatment and neither of age nor ELS treatment x age interaction (Suppl. table 4).

Hence, while ELS leads to an age-independent increase in the number of ΔFosB^+^ neurons in the BLAa in males, in females ELS leads to an age-dependent increase in the number of ΔFosB^+^ neurons in the BLAa specifically during pre-juvenile development. In both males and females the effect was similar in either hemisphere and the intensity of the signal did not change neither with age nor with ELS treatment. These data further support the results from the *in vivo* electrophysiological recordings in the BLAa, and strongly indicate persistent activation of a population of BLAa neurons by ELS in both males and females during pre-juvenile development.

### 3.7 ELS persistently activates non-GABAergic neurons in pre-juvenile BLAa

It has previously been shown that GABAergic interneurons in the BLA, particularly parvalbumin (PV) interneurons, are highly susceptible to ELS, as they show precocious maturation resulting in increased numbers during pre-juvenile development following ELS ^12, 44^. Moreover, their synaptic and structural input is also affected by ELS, leading to altered feed-forward inhibition in the developing amygdala ^26^. To investigate whether those BLAa neurons showing increased activity after ELS as determined by ΔFosB^+^ immunohistochemistry are also GABAergic, we performed triple immunohistochemistry against FosB, PanFosB and the 67 dDa isoform of the GABA synthesizing enzyme glutamate decarboxylase (GAD67) (Fig. 6) in P18 control and ELS mice. As above (see Fig. 5), PanFosB^+^ neurons that did not co-localize with FosB were identified as ΔFosB^+^. To limit the number of animals needed, we conducted these experiments with remaining tissue generated for the experiments described in Figure 5. The data show a similar trend of increased ΔFosB^+^ neurons after ELS (Fig. 6Bi), but did not quite reach statistical significance. However, the number of GAD67^+^ neurons was similar in all four experimental groups (Fig. 6Bii) as was the intensity of the staining which was clearly distinct from background (Suppl. Fig. 5). Interestingly, in neither control nor ELS males, ΔFosB^+^ neurons co-localized with GAD67^+^ neurons at all (0%, Fig. 6A(i), B(iii)), suggesting that they are rather glutamatergic projection neurons. In females though, ΔFosB^+^ neurons co-localized with GAD67^+^ neurons in a low percentage of cells in controls (4 ± 2.72 %) which was similar after ELS (5 ± 5.42 %) (Fig. 6Aii, Biii). There was high variability between mice as half of the controls and 3/4 ELS females showed no co-localization, while 7% or 11% of co-localization was observed in the remaining two control females vs 22% of co-localization in the remaining ELS female.

**Figure 6.**
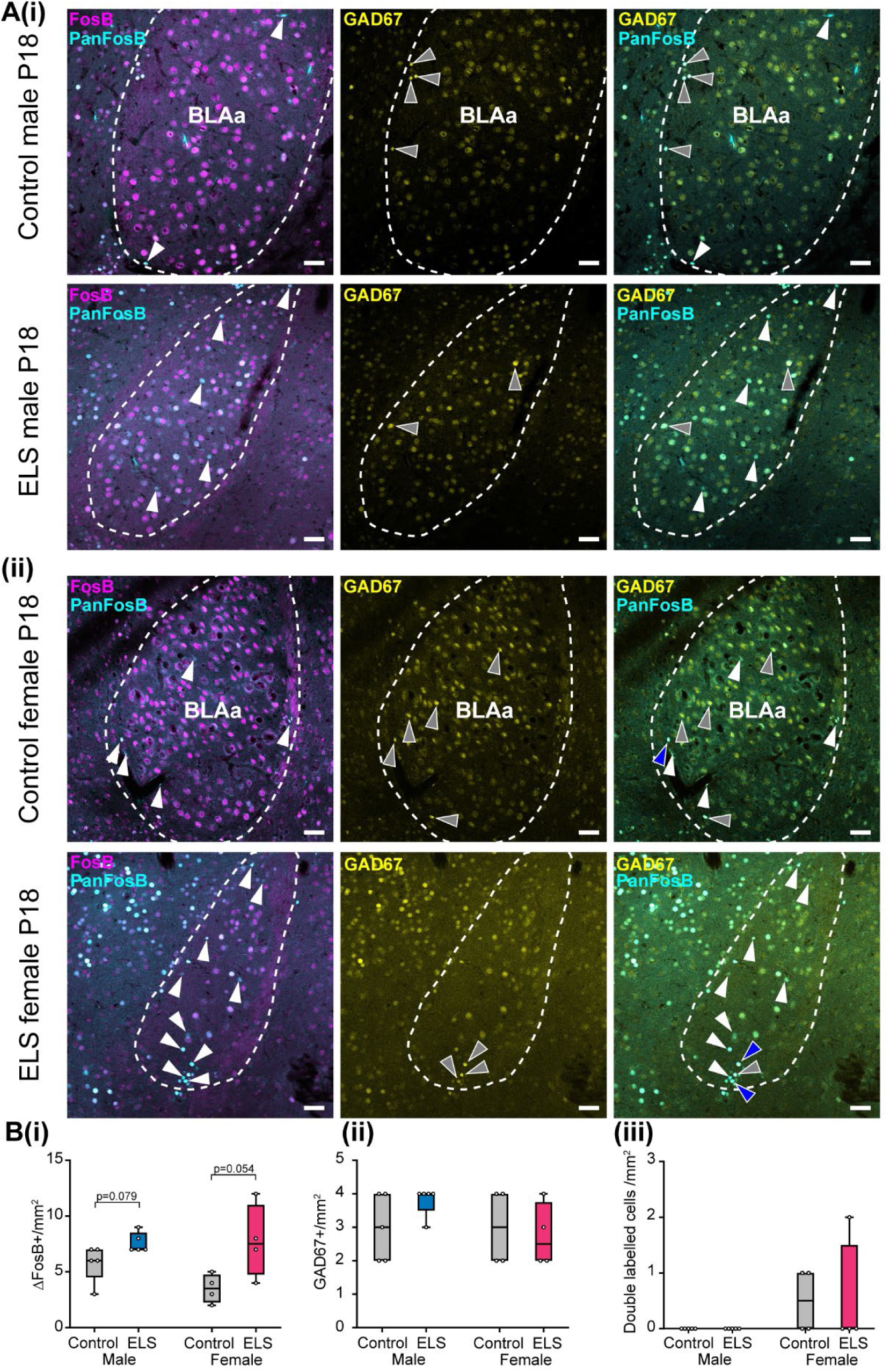
ELS does not cause chronic activation of GABAergic pre-juvenile BLA neurons. **A,** **(i)** Representative photomicrograph images showing neurons immunopositive (+) for FosB (magenta, left), PanFosB (cyan, left) and GAD67 (yellow, center) in the BLAa of a control (top row) and ELS (bottom row) male P18 mouse. The merged images on the left show ΔFosB^+^ cells (white arrowheads), i.e. those PanFosB^+^ cells which do not colocalize with FosB. The merged image on the right shows that those ΔFosB^+^ cells (white arrowheads) do not colocalize with GAD67 (grey arrowheads) in the BLA. **(ii)** Same as (i) for a control and ELS female mouse. Note that while in males ΔFosB^+^ cells never colocalized with GAD67^+^ cells, in both control and ELS females there was some colocalization with GAD67^+^ cells (blue arrowheads). Scale bars: 50 µm. **B,** Box and whisker plots showing the number of ΔFosB^+^ cells/mm^2^ (**i**, left), the number of GAD67^+^ cells/mm^2^ (**ii**, center) and double labelled ΔFosB^+^ and GAD67^+^ cells/mm^2^ (**iii**, right) in the BLAa in control (grey) and ELS (blue: male, magenta: female) mice at P18 (males: N = 5 mice per group, females: N = 4 mice per group). Note the trend towards an increase in ΔFosB^+^ cells after ELS in both sexes, which was close to statistical significance (Mann-Whitney test or paired t-test, respectively). Note that while the number of GAD67^+^ cells is similar in all groups of mice, in control and ELS males GAD67^+^ cells did never colocalize with ΔFosB^+^ cells. In both control and ELS females though, colocalization of ΔFosB^+^ and GAD67^+^ cells was present.

Hence, these data strongly suggest that ELS leads to a persistent activation of predominantly BLAa principal neurons rather than GABAergic interneurons. Whether in females ELS also leads to an activation of GABAergic neurons in the BLAa still has to be determined in a larger dataset.

### 3.8 ELS leads to a lower firing activity of neurons in the prefrontal cortex of pre-juvenile males

To get a better understanding of how ELS affects the development of the pre-juvenile mPFC, we sorted cortical MUA into single units and analysed their firing activity. We focussed our analysis on superficial layers II/III within PL due to its strong connectivity with BLAa ^45, 46^. In control males, we obtained n = 54 units from N = 11 pups and in ELS males n = 90 units from N = 13 pups. Baseline firing rates in control PL were low (1.45 (1.68) Hz) and Fano factor was 1.80 (1.27) indicating high variability in the firing activity (Fig. 7A). Interestingly, baseline firing rates in the PL of ELS mice were significantly lower than in controls (0.89 (1.37) Hz, U = 1902, p = 0.029, Mann-Whitney test) but the Fano factor was similar (1.76 (0.96)). We also analysed the spike waveforms in both groups (Fig. 7B). Both valley-to-peak time (ms) and FWHM were similar between control and ELS males. The decrease in neuronal firing activity within the PL of ELS males may be due to a change in the excitation/inhibition balance or an increased feed-forward inhibition. We relied on the shape of the extracellular waveform to sort all recorded PL units into putative regular spiking excitatory vs putative fast-spiking inhibitory units as previously shown for spiking activity in the pre-juvenile PFC ^32^ (Suppl. Fig. 6Ai). Interestingly, the ratio of putative inhibitory vs excitatory neurons in the PL was significantly different after ELS (p = 0.0531, Fisher’s test). In control males, only one unit (1/54, 2%) was classified as fast-spiking while in ELS males 10 units (10/90, 11%) were classified as fast-spiking. Increased population activity of inhibitory fast-spiking interneurons may contribute to lower baseline firing rates within PL in ELS males. However, immunolabelling against parvalbumin (PV) and subsequent quantification of the density of PV-positive (PV+) neurons in P18 males did not reveal any signifcant differences between control and ELS mice (Suppl. Fig. 8). Thus, it seems that ELS does not lead to higher density of PV+-neurons in BLAa during pre-juvenile development, but rather to an increase in activity of a similar number of PV+ interneurons compared to controls. Moreover, when analysing firing activity within layers V/VI of the IL cortex, which also shows strong connectivity to the BLA ^45, 46^, we also found a decrease in the main firing rates in ELS males (Suppl. Fig. 6B, U = 5527, p = 0.0081, Mann-Whitney test). While the ratio of putative excitatory vs inhibitory units was not different between control and ELS mice (Suppl. Fig. 6D), spikes recorded in the IL of ELS mice had a significantly shorter half-width than in controls (Suppl. Fig. 6C, U = 5328, p = 0.0025, Mann-Whitney test), which may also indicate that more putative inhibitory units with faster action potentials are active after ELS in the IL in males.

**Figure 7.**
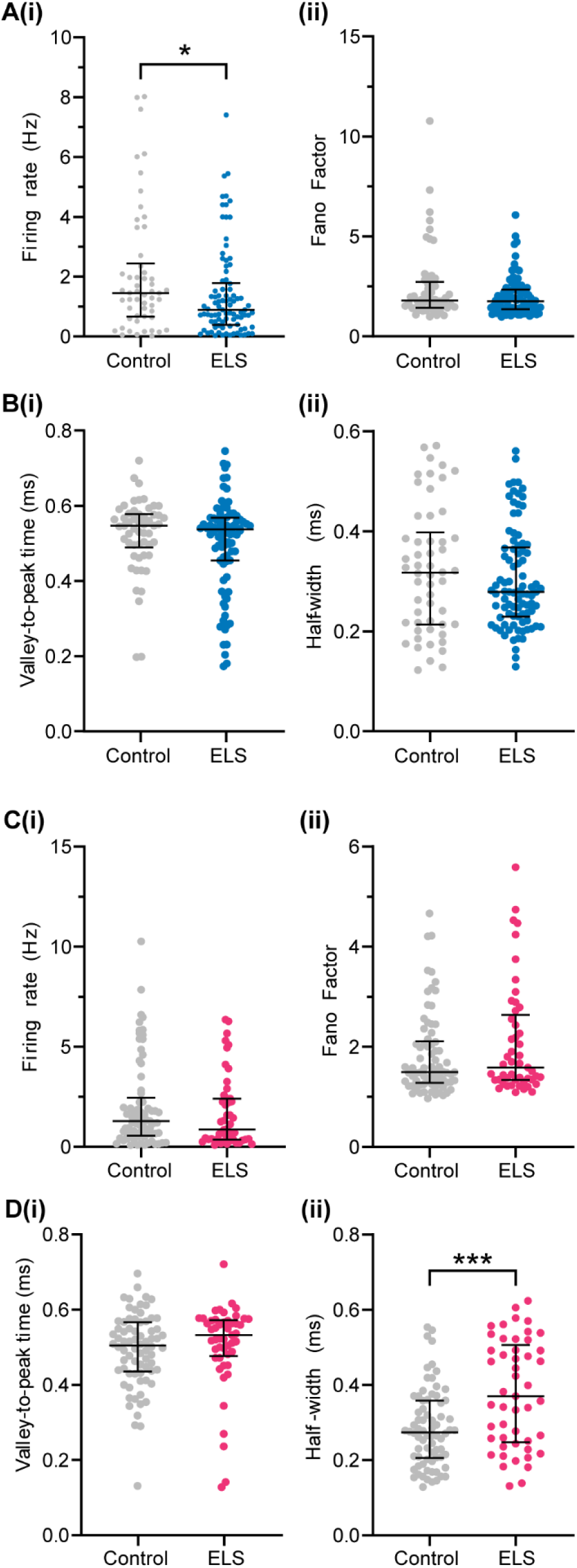
ELS decreases the firing activity of neurons in layers II/III of the prelimbic cortex in pre-juvenile males only. A, **(i)** Distribution and median (IQR) of instant firing rates (Hz) of single units in PL in control (grey, n = 54 units from N = 12 pups) and ELS (blue, n = 90 units from N = 13 pups) male pre-juvenile mice. Note the significant decrease in firing rates after ELS. **(ii)**, Same as A (i) for Fano factor. **B, (i)** Distribution and median (IQR) of the mean valley-to-peak time (ms) corresponding to the mAHP of the extracellular action potential of single units in the PL in control (grey, n = 54 units) and ELS (blue, n = 90 units) male pre-juvenile mice. **(ii)** Same as B (i) for the full width at half maximum (ms) of the extracellular action potential waveform. **C, (i)** Distribution and median (IQR) of instant firing rates (Hz) of single units in PL in control (grey, n = 74 units from N = 13 pups) and ELS (magenta, n = 49 units from N = 10 pups) female pre-juvenile mice. **(ii)**, Same as C (i) for Fano factor. **D, (i)** Distribution and median (IQR) of the mean valley-to-peak time (ms) of single units in the PL in control (grey, n = 74 units) and ELS (magenta, n = 49 units) female pre-juvenile mice. **(ii)** Same as D (i) for the full width at half maximum (ms) of the extracellular action potential waveform. Note the significant increase in half-width after ELS. *p < 0.05, ***p < 0.001, Mann-Whitney test.

In females, after spike sorting we obtained n = 73 units in N = 13 pups in control and n = 49 units from N = 10 ELS pups. Similar to males, firing rates of PL units in female control mice were low (1.28 (1.74) Hz) and Fano factor was 1.50 (0.81), the latter indicating a high variability in firing activity over time (Fig. 7C). Both the firing rate (0.87 (2.03) Hz) of PL units as well as the Fano factor (1.59 (1.26)) were similar in ELS mice. Furthermore, the ratio of putative regular spiking excitatory neurons vs putative fast-spiking inhibitory neurons was also similar between the groups (Suppl. Fig. 6Aii). When analysing the valley-to-peak time (ms) of the PL spike waveforms, their length was similar in both control and ELS females (Fig. 7D). However, the FWHM of the initial negative peak of the extracellular action potential was significantly longer in ELS females than in control females (control: 0.27 (0.15) ms, ELS: 0.37 (0.25) ms, U = 1136, p = 0.0004, Mann-Whitney test).

For adolescent mice, after spike sorting we obtained n = 12 single units from N = 6 control males and n = 23 units from N = 9 ELS males as well as n = 52 single units from N = 14 control females and n = 30 single untis from N = 14 ELS females. Our analysis revealed no significant differences in either sex for any of the parameters analysed between control and ELS mice (Suppl. Fig. 7 A-D). We observed a trend towards a decrease in mean firing rates in ELS males, which appears to be in line with our observation in pre-juvenile males. However, due to the low number of units in controls, this analysis would have to be confirmed in a larger dataset.

These data suggest that during pre-juvenile development, ELS leads to an overall decrease in firing activity in the PL in males. Furthermore, increased population activity of inhibitory fast-spiking interneurons may contribute to hypoactivity within mPFC networks after ELS in pre-juvenile males. During adolescent development, ELS males showed a trend towards lower mean firing rates in PL which would have to be confirmed in a larger data set.

In females, neither the firing rate nor the firing pattern were affected in PL by ELS, and also no changes in the mAHP of the spikes were detected during either developmental time point.

### 3.9 ELS disrupts spike-LFP synchronization both locally and between the BLAa and the mPFC

To further characterize the effect of ELS on long-range interactions between mPFC and BLAa, we sought to estimate the phase synchronization between spikes and LFP, both within and across PFC and BLAa. We focussed on the superficial layers of the PL and the LFP in the 3-5 Hz low-theta band in pre-juvenile mice, as impairments in coherence were observed in this frequency band and this subregion of the mPFC. In the BLAa of control males, local spike firing occurred preferentially on the descending phase of the theta cycle (Fig. 8Ai), with the vast majority of units (31/36, 86%) being locked to the local theta rhythm. BLAa units in ELS males showed a similar preferred phase angle to the local theta rhythm and a comparable locking strength as in controls, as measured by the length of the mean resulting vector. The vast majority (28/29, 97%) of BLAa units in ELS males was also locked to the local theta rhythm. BLAa units in control females also occurred at a similar preferred phase angle to their male counterpart (Fig. 8Aii), with almost every unit being locked to the descending phase of the local LFP (33/35, 94%). ELS females also showed a high proportion of locked BLAa units (19/22, 86%) as well as a similar locking strength to control females, but the preferred phase angle differed between the two groups (Watson-Williams test, p = 0.0062). While this difference in phase angle was consistent and statistically significant, it was only ∼20° in absolute terms, meaning that the spikes were still functionally locked to a similar cycle phase in both groups.

**Figure 8.**
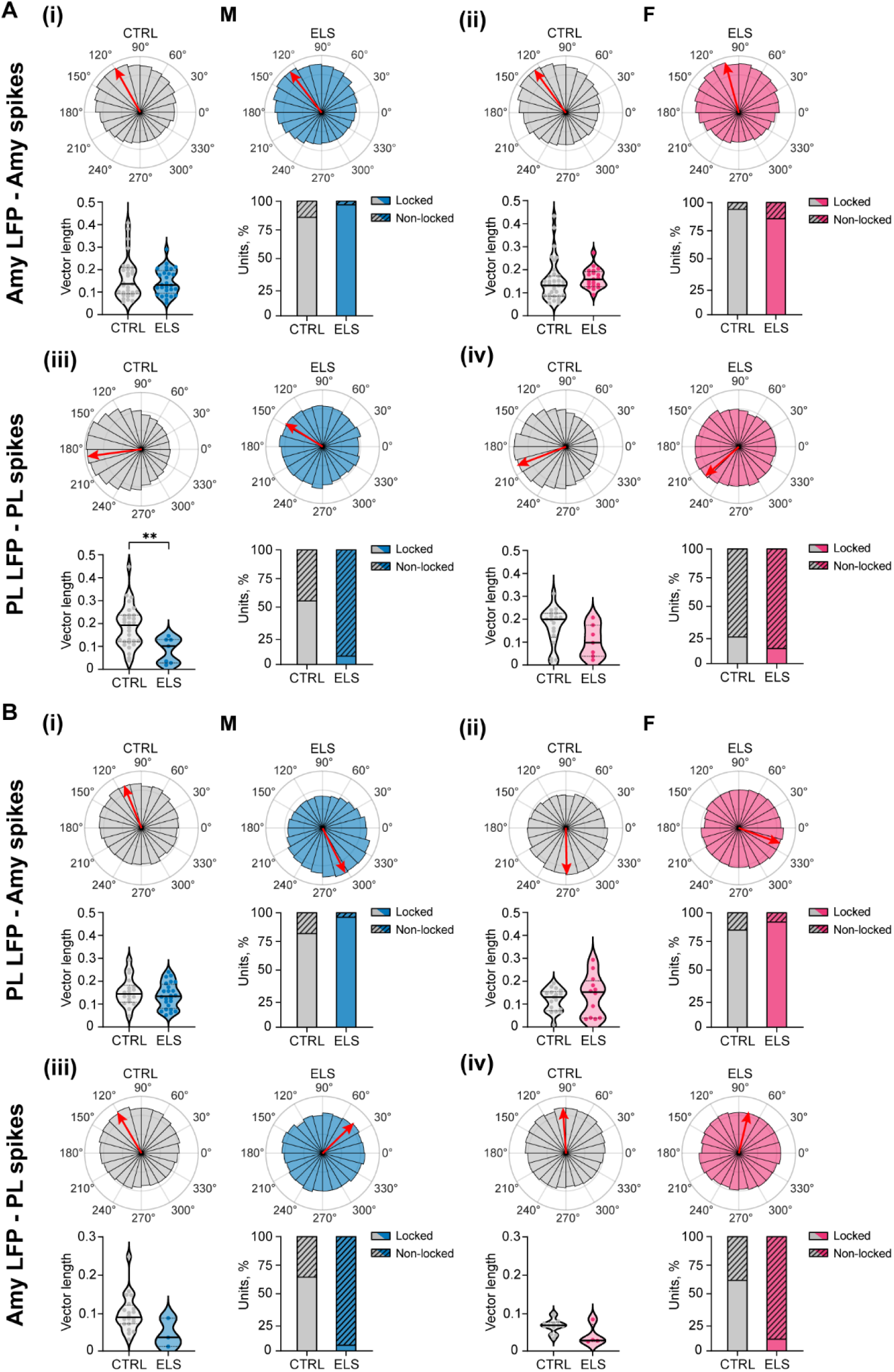
ELS affects the locking of spikes to the low-theta rhythm within prefrontal-amygdala networks. A, **(i)** Top: Polar plots showing the probability distribution of the preferred BLAa spike phase locked to the 3-5 Hz LFP within the BLAa itself, for control (left, grey) and ELS (right, blue) male pre-juvenile mice. Red arrows show the vector mean phase for each group, and their length was scaled to match the height of the tallest bin. 0° corresponds to the peak of the theta cycle, and 180° to its through. Bottom: comparison of the mean vector length for each unit (left) and the relative proportion of locked (solid colors) and non-locked (striped pattern) units within each group (right). **(ii)** Same as A (i) for female control (left, grey) and ELS (right, magenta) pre-juvenile mice. **(iii)** Same as A (i) for PL spikes phase locked to the 3-5 Hz LFP within the PL itself, for control (left, grey) and ELS (right, blue) pre-juvenile male mice. **(iv)** Same as A (iii) for female control (left, grey) and ELS (right, magenta) pre-juvenile mice. **B, (i)** Top: Polar plots showing the probability distribution of the preferred BLAa spike phase locked to the 3-5 Hz LFP of the PL, for control (left, grey) and ELS (right, blue) male pre-juvenile mice. Red arrows show the vector mean phase for each group, and their length was scaled to match the height of the tallest bin. Bottom: comparison of the mean vector length for each unit (left) and the relative proportion of locked (solid colors) and non-locked (striped pattern) units within each group (right). **(ii)** Same as B (i) for female control (left, grey) and ELS (right, magenta) pre-juvenile mice. **(iii)** Same as B (i) for PL spikes phase locked to the 3-5 Hz LFP of the BLAa, for control (left, grey) and ELS (right, blue) pre-juvenile male mice. **(iv)** Same as B (iii) for female control (left, grey) and ELS (right, magenta) pre-juvenile mice. **p < 0.01, Mann-Whitney test.

Within the PL, both in control and ELS males, spikes occurred preferentially at the trough of the local theta cycle (Fig. 8Aiii). However, the locking strength, i.e. length of the mean resulting vector, was significantly smaller in ELS males (Mann-Whitney test, p = 0.009) and also the proportion of PL units locked to the local theta rhythm was much lower than in controls (control: 30/54, 56%, ELS: 7/90, 8%, Fisheŕs test, p < 0.0001), thus indicating a desynchronization between the spiking activity and the local theta rhythm. Interestingly, none of the putative fast-spiking interneurons in PL in either treatment group showed significant locking, suggesting that locked units are mainly principal neurons. Both in control and ELS females, PL units also preferentially fired at the trough of the local theta cycle (Fig. 8Aiv). In both groups, the number of significantly locked units was low (control: 18/74, 24%, ELS: 7/49, 14%) and not different between the groups. However, as in males, there was a trend towards a decreased vector length (i.e. locking strength) in ELS females vs controls, but this did not quite reach significance (Mann-Whitney test, p = 0.097) probably due to the low number of locked units. Overall, while ELS caused prominent desynchronization between the firing of PL units and the local theta rhythm in males, this effect was much weaker in females, suggesting that developing prefrontal networks in males are particularly vulnerable to ELS.

When analysing long-range spike-field interactions, in control males a high percentage of BLAa units (18/22, 82%) was locked to the theta rhythm in PL, with spikes preferentially occurring on the descending phase of the PL theta cycle (Fig. 8Bi). In ELS males, a similar proportion of BLAa units was locked to the PL theta rhythm (24/25, 96%), and also the vector length was similar. However, their preferred phase angle was dramatically different (Watson-Williams test, p < 0.0001), displaying a ∼180° inversion, with BLAa spikes being locked to the ascending phase of the PL theta cycle. In control females, also a high percentage of BLAa units (17/20, 85%) was locked to the PL theta rhythm, but in stark contrast to control males, spikes occurred preferentially on the ascending phase of the PL theta cycle (Fig. 8Bii). In ELS females, the proportion of locked BLAa units as well as the locking strength to PL theta was similar to control females. However, as for ELS males, ELS also affected the preferred phase angle in females (Watson-Williams test, p = 0.039) with spikes occurring preferentially later in the cycle closer to the peak than in control females, although the difference in phase angle was not as dramatic as in males.

When analysing phaselocking of PL units to the BLAa theta rhythm, in control males the majority of PL units (20/31, 61%) was locked to the BLAa theta rhythm with spikes occurring at the descending phase of the theta cycle (Fig. 8Biii). In ELS males, very few PL units (3/55, 6%) were locked to the BLAa theta rhythm, a significantly lower proportion compared to controls (Fisheŕs test, p < 0.0001) and none of them were putative fast-spiking interneurons. Those few units that were locked displayed also a different preferred phase angle (Watson-Williams test, p = 0.012) occurring earlier in the theta cycle than units in controls. In control females, similar to males, the majority of PL units was locked to the BLAa theta rhythm (13/21, 62%) with a preferred phase angle on the descending phase of the BLAa theta cycle (Fig. 8Biv). As in ELS males, also in ELS females very few PL units (4/37, 11%) were locked to the BLAa theta rhythm, significantly less than in control females (Fisheŕs test, p < 0.0001). However, their preferred phase angle was similar as in controls. Neither in ELS males nor females the vector length was significantly different to controls, although this was difficult to assess due to the low number of locked PL units in ELS pups.

Overall, spike-LFP phaselocking analysis showed a prominent effect of ELS on the entrainment of spiking activity in the PL, predominantly in males, to both the local and the BLAa theta rhythm, resulting in a lower proportion of locked units, weaker locking strength, and a shift in the preferred phase angle. ELS females had a milder phenotype limited to long-range outcomes. Interestingly, ELS also led to a major change in the preferred phase angle of BLAa units locked to PL theta. Together with the effect of ELS on the entrainment of PL spikes, this suggests that ELS results in prominent changes in how synaptic inputs are integrated in the PL and may lead to an abnormal output from the developing PL.

## 4. Discussion

The present study shows that ELS causes major dysfunction within prefrontal-amygdala networks *in vivo* during pre-juvenile development in a sex-dependent manner. Hence, this developmental period may present a “window of opportunity” during which an abnormal developmental trajectory inflicted by ELS may still be rescued. We found evidence of changes in local neuronal firing activity after ELS, such as increased firing activity within BLA but decreased firing activity within mPFC. Long-range functional interactions were also affected after ELS leading to exaggerated oscillatory theta coupling as well as deficits in the reciprocal theta entrainment of spikes. These early impairments in prefrontal-amygdala circuit function may constitute an altered developmental trajectory towards an increased susceptibility to psychiatric diseases later in life.

### 4.1 Oscillatory network activity within developing mPFC or amygdala is not affected by ELS

Temporal coordination through neuronal synchrony as seen during oscillatory network activity is critical for normal adult brain function ^47^. Moreover, impaired oscillatory network activity has been found during postnatal development in rodent models of psychiatric disorders, and may even constitute an early biomarker of later disease, for review see ^48^. To address if ELS also impacts neuronal synchrony within developing prefrontal-amygdala networks, we performed LFP recordings within pre-juvenile and adolescent prelimbic, infralimbic, and BLAa networks of control and ELS mice under light urethane anaesthesia, and we analysed the PSD of frequencies spanning the 0.1-100 Hz range. In addition to the anaesthesia induced slow-wave activity, we found predominant activity in the low-theta range (3-5 Hz) within mPFC networks at both ages. This is in line with recent reports in awake mice that also describe predominant network activity in the low-theta range during resting at similar ages ^32^. Although we did not quantify this, our data also indicate an increase in high-theta (6-12 Hz) as well as in gamma power (12-100 Hz) in the mPFC from pre-juvenile to adolescent development (see Fig. 1C) similar to what has been observed in awake mice in the mPFC ^32^. This suggests that the gross developmental changes in prefrontal network activity are also visible and preserved under anaesthesia and further justifies the focus of our analysis on the low-theta band as the dominating network activity in pre-juvenile mice. However, we did not find any effect of ELS on PSD neither in the mPFC nor in the BLAa. While this could be due to the technical limitation of performing all recordings under anaesthesia, a recent preprint on the effect of ELS on low-theta power (2-5 Hz) in mPFC or amygdala of adult awake behaving mice did not find any significant changes neither during resting nor following fear conditioning (during extinction or recall) in both sexes ^49^. These findings strengthen our conclusion that ELS does not impact the generation of synchronous network activity within the developing mPFC nor BLAa per se particularly in the low-theta range.

### 4.2 ELS leads to sex-specific prefrontal-amygdala theta-hypercoupling during pre-juvenile development

Impaired prefrontal-amygdala long-range interactions are a major disease hallmark after ELS, and are also present in subjects at risk ^4, 50^. Previous studies showed impairments in prefrontal-amygdala functional connectivity obtained with fMRI both during development and in adulthood in rodent models of ELS ^15, 23–25^. However, direct electrophysiological measurements of neuronal network activity in these models remain sparse ^26^. We analysed coherence as a quantitative measure of oscillatory coupling within prelimbic- or infralimbic-BLAa networks. We focussed on the low-theta band (3-5 Hz), as impairments in long-range low-theta synchrony were present in our previous study in pre-juvenile ELS rat pups ^26^. Moreover, theta oscillations within this frequency band are known to support long-range coordination such as fear learning and extinction, and they are also a substrate of innate anxiety ^51–53^. Our findings highlight an increased coupling between the BLAa and the PL in layers II/III in the low-theta band which was specific to pre-juvenile males, while females had a milder phenotype showing only a trend towards theta hypercoupling. This is in line with previous reports also showing that early impairments in functional interactions after ELS are limited to males ^23, 25, 26^. No differences in theta coupling were observed between control and ELS mice during adolescence. However, when analysing the increase in prefrontal-amygdala low-theta coherence from pre-juvenile development to adolescence, we found that this increase was significantly lower in both ELS males and females, which may indicate a precocious development of coupling strength after ELS. Previous studies showed a decrease in functional interactions within prefrontal-amygdala networks after ELS in pre-juvenile rats ^23, 25, 26^ using either MS or LBN as a stress model. Differences between species, in the stress paradigm, and particularly in the timing of stress exposure may underlie the different outcomes in the valence of the effect between those of the previous and our study. However, in line with those previous studies we also found that the effect of ELS on developing prefrontal-amygdala networks is not linear, i.e. it does not simply increase over time, but rather it is confined to certain developmental windows, often during juvenile development, which appear to be more susceptible to stress^6, 25, 40^. Accordingly, early theta-hypercoupling during pre-juvenile development may either correspond to the previously described accelerated development of prefrontal amygdala circuits or may constitute an impairment confined to pre-juvenile development which seems to be particularly vulnerable to ELS. Further studies are necessary to corroborate this finding and its meaning. Interestingly, similar cumulative ELS (LBN+MS) in mice also resulted in increased resting state functional connectivity ^15^ as well as anatomical hyperconnectivity ^24^ during adulthood in males only, which strongly correlated with an anxiety phenotype ^15^. Behavioural characterization of our LBN+MS ELS model during adulthood revealed increased anxiety-like behaviours in ELS males and strong deficits in classical fear conditioning in both ELS males and females^27^. Future studies are needed to confirm if hypercoupling within prefrontal-amygdala networks also emerges during adulthood in our ELS model and contributes or underlies an anxiety phenotype.

### 4.3 ELS leads to chronic increase in firing activity of a subpopulation of putative (non-GABAergic) principal neurons in the BLAa during pre-juvenile development and to a more widespread increase in mean firing rates during adolescence

Analysis of *in vivo* spike firing activity as well as immunohistochemistry against ΔFosB in pre-juvenile BLAa suggest that ELS leads to an increase in firing activity of a subpopulation of BLAa neurons already during brain development. Increased activity of amygdala neurons after ELS has been previously described as an increase in the number of cFos positive neurons in response to a stressful stimulus ^16, 17^. Furthermore, amygdala hyper-reactivity is one of the most consistent findings after ELS in humans ^4^. Here we extend these findings to ELS models, and we show for the first time that ELS also leads to higher baseline firing rates in a subpopulation of BLAa neurons *in vivo* in either sex. The shorter valley-to-peak time of the spike waveforms in ELS males vs controls (corresponding to the medium afterhyperpolarization, mAHP) is a further indicator of increased excitability of BLAa neurons after ELS. Chronic stress during either adolescence or adulthood has previously been shown to lead to a decrease in the amplitude of the mAHP, as well as to an increase in the intrinsic excitability of BLA pyramidal neurons in *in vitro* brain slice recordings in adult rats or mice ^20, 54^. These changes were due to the loss of currents from small-conductance calcium-activated potassium channels containing subunit 2 (SK2), e.g. by reduced expression after ELS ^20, 54^. Furthermore, altered excitatory neuropeptide release may also contribute to the reduced mAHP and potentially increased neuronal excitability in BLAa after ELS. For example, corticotropin-releasing hormone (CRH) is expressed in a significant population of BLA neurons^55^ and has been shown to positively modulate neuronal excitability through effects on potassium channels and is also known to lead to a decrease in AHP decay time^56^.

Further experiments are needed to reveal if downregulation of SK2 channels and or altered CRH release after ELS in male pre-juvenile BLAa also underlies the change in length of the mAHP in our ELS model, which may promote increased baseline firing rates at least in a subpopulation of BLAa neurons. Of note, we did not find any changes in the valley-to-peak time of BLAa spikes in female pre-juvenile pups, suggesting that the increase in baseline firing of a subpopulation of BLAa units in females may have a different underlying mechanism than in males. Our analysis of BLAa neuronal activity during adolescence revealed an even stronger and more widespread increase in mean firing rates in ELS males as well as impaired firing patterns compared to controls, suggesting that the phenotype further increased from pre-juvenile to adolescent development. The data in adolescent females point to a similar phenotype as in males, but would have to be confirmed in a larger dataset. The decrease in BLAa spike FWHM in both ELS males and females suggests either altered expression of ion channels such a sodium or calcium channels after ELS or may result from the chronic increase in activity of these neurons. Most likely these neurons with higher baseline firing activity mostly comprise BLAa principal neurons, as we did not find any indication from correlation or clustering analysis for increased activation of FSIs. Interestingly, the decrease in valley-to-peak time corresponding to mAHP did no longer persist in ELS males during adolescence, but was present in ELS females instead, suggesting increased excitability of BLAa neurons in adolescent females and combined with the decreased half-width of the spike and overall shorter spike duration.

In line with the effect of ELS on pre-juvenile spike firing in BLAa, our immunohistochemistry data showing an increase in ΔFosB^+^ neurons in the BLAa after ELS in both sexes provides further evidence that ELS leads to a persistent increase in neuronal firing activity of a subpopulation of neurons within pre-juvenile BLAa. The Δ splice variant of the immediate early gene FosB is a suitable marker for the effect of chronic stress on neuronal activity ^43^ and has previously been shown to also respond to chronic stress within the developing amygdala in rats ^57^. Due to the unavailability of commercial antibodies against ΔFosB, double staining against PanFosB and FosB to identify neurons expressing ΔFosB has been precedently established as an alternative method ^58^ and has also worked well in our study. Interestingly, while in males the increase in ΔFosB^+^ neurons was independent of age and already present right after the end of the chronic stress period, in females it only emerged at P18, which may point to a different underlying mechanism for the increase in neuronal activity than in males. Future experiments are needed to reveal the identity of the activated neurons, and whether they share a specific projection target or they receive a distinct set of inputs. Our triple immunohistochemical staining against ΔFosB and GAD67 already revealed that ΔFosB^+^ neurons were GAD67 immunonegative in pre-juvenile males, while in pre-juvenile females some double positive neurons were encountered. Hence, the vast majority of those chronically activated neurons in the pre-juvenile BLAa do not seem to be GABAergic interneurons. It has previously been shown that chronic stress in adulthood leads to the activation of neurons that project to the ventral hippocampus (vHipp), which is associated with a selective decrease in SK2 channel currents as well as an increased intrinsic excitability in these cells ^54^. Moreover, overexpression of SK2 channels in vHipp-projecting BLA neurons rescued the anxiety phenotype after ELS. Future research will show whether similar target specificity also applies to developing BLA neurons activated by ELS.

### 4.4 ELS leads to a decrease in firing rates *in vivo* in the pre-juvenile PL in males

In contrast to the BLAa, where ELS resulted in an increased activity of a subpopulation of neurons, its effect on baseline firing activity within layer II/III of the pre-juvenile PL was broadly inhibitory. In males, mean baseline firing rates of PL units were decreased in ELS pups compared to controls. Our results are in line with previous data claiming that ELS leads to inhibition of the mPFC. It has been shown that ELS leads to a decrease in synaptic spine density on pyramidal cells within both the superficial and the deep layers of the adult PL, which is associated with decreased glutamatergic transmission in the PL after ELS ^59^. Accordingly, chemogenetic activation of PL pyramidal neurons in ELS pups rescued deficits in working memory performance. Furthermore, in line with the hypothesis that ELS leads to an accelerated development, the GABA switch from excitatory to inhibitory and also the downregulation of the excitation/inhibition balance happens earlier after ELS, leading to an attenuated excitatory tone within the developing mPFC ^60, 61^. Finally, chemogenetic postnatal inhibition of the mPFC between P2-17 resulted in similar cognitive deficits in adulthood as those observed following ELS, and chemogenetic activation of the mPFC during ELS was able to prevent cognitive deficits in adulthood ^62^. Our data contribute to this literature and show, for the first time, that ELS also results in decreased firing rates in the mPFC *in vivo* already during pre-juvenile development. This decrease in firing rates may hamper the ability of the mPFC to control subcortical networks such as the BLAa, which may further result in increased neuronal activity therein, and may promote disease symptoms after ELS. Interestingly, we did not find an effect of ELS on PL mean firing rates in pre-juvenile females, suggesting a milder phenotype in females. However, the increase in half-width after ELS also suggests that ELS may alter the expression or function of, e.g. K^+^-channels on PL pyramidal neurons, thus affecting the repolarization phase of the action potential. Future studies are necessary to further explore the functional meaning of such changes for PL network activity in pre-juvenile females. Interestingly, none of the changes in spike firing observed during pre-juvenile development persisted during adolescence, suggesting that pre-juvenile PL is particularly vulnerale to ELS. However, a trend towards decreased firing rates in PL of adolescent ELS males compared to controls may point to an inhibitory effect of ELS on PL network activity also during adolescence. However, this would have to be confirmed in a larger dataset.

### 4.5 ELS differentially affects spike-LFP synchrony in males and females

The Amy and the PFC are tightly interconnected in a bidirectional circuit, with the Amy sending salience information to the PFC, and in turn receiving top-down regulatory inputs ^63^. While our coherence analysis revealed abnormal hypercoupling within prelimbic-BLAa networks after ELS in pre-juvenile males, this analysis did not provide any information regarding the effect of ELS on directionality within this network, i.e. whether interactions from Amy→mPFC or from mPFC→Amy were affected. Hence, we analysed the phaselocking of spikes in the PL and in the BLAa to either the local low-theta rhythm or long-range to the low-theta rhythm in the BLAa or in the PL, respectively. Phase synchronization supports functional communication between distant areas through coordinated spiking activity at specific oscillatory phases ^64–66^. While the entrainment of BLAa spikes to the local theta rhythm was not affected by ELS, the entrainment of PL spikes to the local theta rhythm was impaired after ELS in pre-juvenile males, resulting in much lower locking strength as well as a much smaller proportion of locked units. Moreover, the ability of the BLAa to entrain PL spikes to the BLAa theta rhythm was also hampered after ELS particularly in males, resulting in only a minority of units displaying significant locking, as well as a change in the phase angle. The dramatic decrease in the ability of both the local theta rhythm and the theta rhythm in the BLAa to entrain PL firing after ELS in males strongly suggest that ELS leads to major changes in the integration of synaptic inputs in the PL likely resulting in a desynchronized output of spike firing. This may be due to changes in the anatomical connectivity as hyperconnectivity between mPFC and amygdala has been found in a similar ELS model in pre-juvenile ^26^ or adult males ^24^. Furthermore, a decrease in synaptic spine density on pyramidal cells within the PL that has been shown to be associated with diminished glutamatergic transmission and a decreased expression of GluR1 and NR1 subunits of AMPA or NMDA receptors after ELS ^59^ may also be a contributing factor. Finally, changes in the excitation/inhibition balance due to the impact of ELS on the development of e.g. PV^+^ interneurons^67^ may also affect the integration of synaptic inputs, leading to poor entrainment of PL units.

Moreover, while PL theta was still able to entrain the firing of BLAa units after ELS in both sexes (i.e. locking strength and percentage of locked units were similar to controls), the phase angle (i.e. the timing within the theta cycle) had dramatically shifted after ELS, particularly in males. This further supports altered synaptic integration within these networks after ELS. Since the precise timing of BLA spikes to PFC theta is critical for information transfer from PFC to BLA (i.e. top-down control) to signal safety during both learned and innate anxiety ^53^, dramatic phase-shifts in locking of BLA spikes to PL theta may also interfere with the ability of PFC to dynamically control activity within BLA networks as well as the anxiety response. Hence, our results suggest that ELS affects interactions in both directions from BLAa to PL as well as top-down from PL to BLAa in pre-juvenile males. In females, the effect of ELS on theta entrainment of spikes within or across PL or BLAa where much milder than in males, in line with an only mild effect of ELS on theta coherence in females. This is in accordance with previous reports that females are less susceptible to ELS than males in rodent models ^19, 24, 26^. Further studies are needed to understand whether abnormal functional interactions are only transient or are also present in adulthood, and if they underlie increased anxiety or abnormal fear learning in this model.

### 4.6 Effect of urethane anaesthesia on neuronal network activity

*In vivo* electrophysiological recordings in young mice bare their own challenges. Head-implants for freely-moving recordings are usually too heavy and may interfere with nursing. Head-fixed recordings lead to a strong acute restrained-stress response which is known to strongly activity BLA in adult rodents and may overlay any chronic changes in activity due to ELS. Therefore, we opted to perform *in vivo* electrophysiological recordings in developing mice under anaesthesia. It has been previously shown that brain activity under urethane anaesthesia closely resembles that under natural sleep. For instance, cyclical alternations of brain state during sleep and the alternations induced by urethane are remarkably similar^37^. This suggests that such a high similarity likely reflects shared physiological mechanisms between sleep and urethane-induced anaesthesia, rendering urethane as an anaesthetic ideally suitable for studies examining cyclical brain rhythms. This also has been confirmed when investigating the effect of urethane on network activity during brain development^68^. Moreover, urethane only minimally alters neuronal firing patterns compared to other anaesthetics^69^ The cortical neuronal network activity under anaesthesia we observed was dominated by low-theta oscillations (3-5 Hz), similar to cortical network activity recorded in head-fixed awake mice at similar ages^32^. Therefore, we believe that the neuronal network activity we recorded under urethane anaesthesia shares similar physiological mechanism with network activity in awake rodents at similar ages during natural sleep and to some extent under quiet resting conditions. However, how our data compares to network activity in awake freely moving animals at similar ages and how ELS may affect neuronal network activity during active behaviours such as exploration, decision making or fear learning has to be investigated in future studies.

### 4.7 Sex differences in the effect of ELS on developing prefrontal-amygdala network activity

It has been widely demonstrated, that ELS seems to have sex-specific effects in rodent models but also in exposed humans^24,70^. Typically, either males show a stronger phenotype than females or ELS seems to affect different behavioural traits. For example, it has been shown that ELS specifically promotes depressive-like behaviours in female rodents ^14^. In line with these previous findings, our data also shows stronger effects of ELS in males than females. We observed an increased in coherence in prelimbic-amygdala networks only in pre-juvenile males but not females and also the effect of ELS on spike firing in either PFC or BLAa seem stronger in males than in females. Similarly, theta entrainment of spikes within prefrontal-amygdala networks was also more strongly affected in males than in females. Several complementary mechanisms may contribute to the enhanced male vulnerability we observed. Males show higher baseline rates of neuronal apoptosis during development^71^, which could create greater susceptibility to stress-induced cellular damage during critical developmental windows. Additionally, sex differences in stress hormone receptor expression and sensitivity may amplify the impact of developmental stress exposure in males compared to females. The timing of our stress protocol (P4-P14) corresponds to a critical time window for development, when sex differences in neural maturation are emerging, potentially resulting in different outcomes despite the stressor being equal. Our electrophysiological findings may reflect this differential developmental trajectory, with males showing more pronounced disruptions in neuronal functional properties following ELS. Future studies are needed to further our understanding on the underlying mechanism leading to distinct outcomes of ELS in males versus females.

## 5. Integrative framework and conclusions

Here we show that ELS leads to theta hypercoupling within prefrontal-amygdala networks during pre-juvenile development only, particularly in male pups (Fig. 9). This early hypercoupling is either the result of precocious development and may be independent of any deficits in directed interactions, or may constitute a developmental impairment limited to pre-juvenile development as particularly vulnerable to ELS. Alternatively, it may be driven by a third region such as the ventral hippocampus, which has strong anatomical connectivity with both the mPFC and the BLA. The emerging theta hypercoupling may constitute a compensatory mechanism to an awry reciprocal entrainment within developing prefrontal-amygdala networks. Furthermore, in both males and females, ELS caused a chronic increase in neuronal spiking activity of a subpopulation of neurons in pre-juvenile BLAa, which were non-GABAergic and hence most likely principal neurons. Moreover, in ELS males this phenotype further increased during adolescence, leading to a prominent increase in mean firing rates in a wider population of BLAa neurons. In contrast, in the mPFC, ELS resulted in a net decrease in neuronal firing activity in male pre-juvenile pups. We also showed that directed interactions by theta entrainment of spikes across prelimbic-amygdala networks are also impacted by ELS in pre-juvenile males, leading to weak locking of PL units to both the local and BLA theta rhythms. Overall, our data suggests that ELS has profound effects on the functional development of mPFC networks particularly during pre-juvenile development, leading to a dampened firing activity, i.e. output which is not synchronized and hard to entrain by subcortical networks such as the BLA, but also affects the timing and integration of synaptic inputs within the mPFC. These impairments may increase susceptibility to disease leading to aberrant development of emotional behaviours or cognitive abilities after ELS. Moreover, early impairments in functional coupling as well as directed interactions may impact fear and safety learning. Theta hypercoupling may compromise the ability of the network to express fine-tuned increases and decreases in prefrontal-amygdala theta synchrony, which are associated with fear acquisition and extinction learning ^53, 65^. Since safety is signalled by the mPFC entraining BLA firing via oscillatory theta band activity ^53^, ELS-evoked major phase shifts in theta entrainment may be disruptive. This could result in fear generalization, as animals that generalize fear show high theta power and synchrony to both aversive and non-aversive cues and lack the prefrontal theta entrainment of BLA networks ^53^. However, also an opposite effect on fear and anxiety behaviours may be possible. Since we observed major deficits in BLA theta entrainment of PL spikes, i.e. disruption of bottom-up communication critical to signal potential threats in the environment, juvenile animals may also become less anxious after ELS.

**Figure 9.**
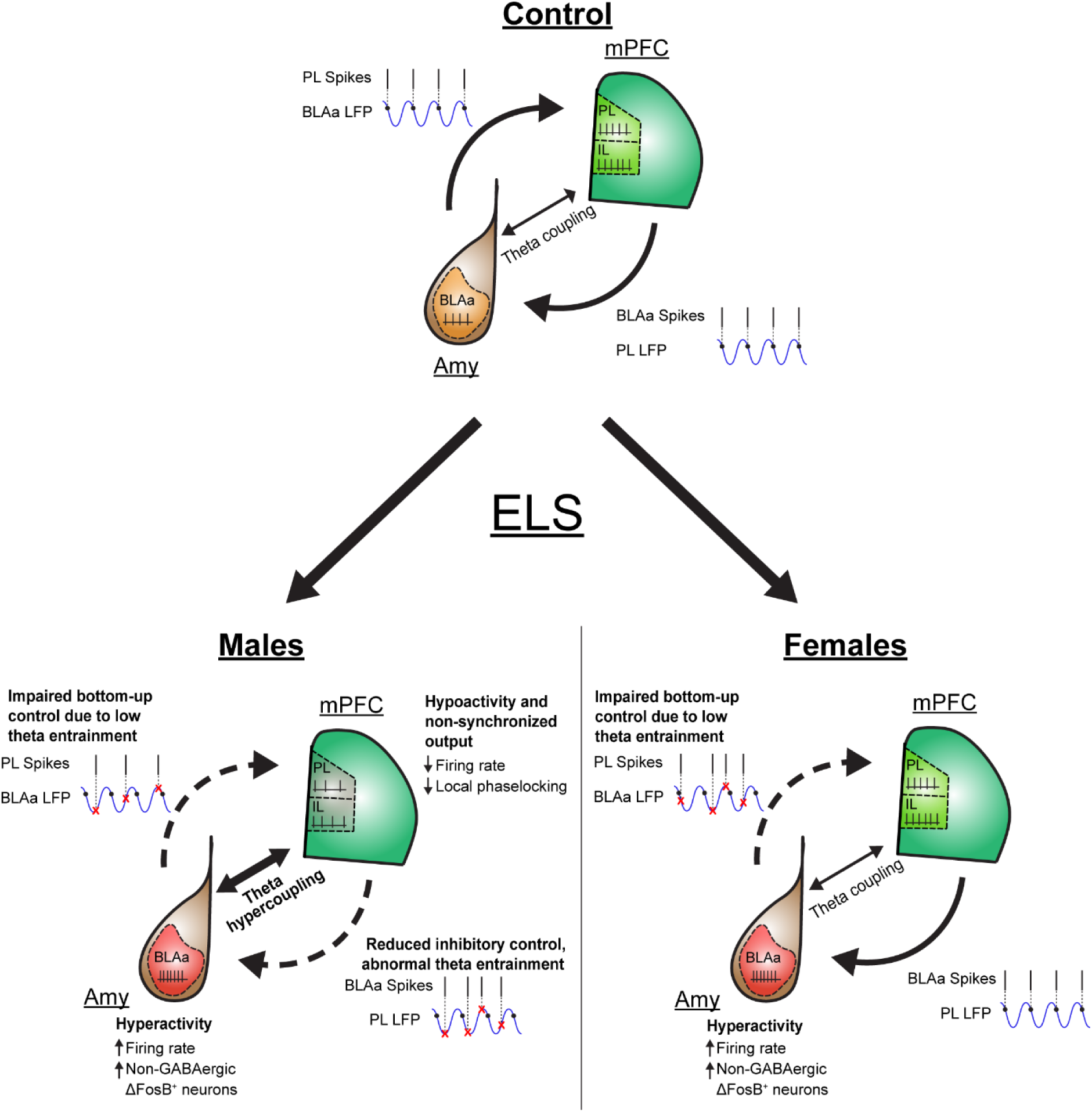
Summary of the sex-dependent effects of ELS on pre-juvenile prefrontal-amygdala networks. Diagram showing baseline neuronal firing activity as well as functional interactions within juvenile prefrontal-amygdala networks in controls (top) and after ELS in male (bottom left) or female (bottom right) pups.

We identified major impairments in prefrontal-amygdala network function following ELS already during pre-juvenile development. Hence, this developmental period may open a window of opportunity during which targeted interventions could prevent the emergence of a disease phenotype during adulthood. Future studies are needed to understand if these impairments within pre-juvenile development lead to persistent changes in the developmental trajectory of prefrontal-amygdala network function, if and how they contribute to a behavioural phenotype, and whether interventions during this developmental window can prevent future disease symptoms.

## Acknowledgments

We thank Assel Kalmenova and Nicolas Vetter for technical help. This study was financially supported by the Research Council of Finland (project grant (decision no 333341) to H.H.

## Author contributions

A.D. and F.V. performed the *in vivo* electrophysiological recordings, F.V. analysed LFP data, A.D. performed spike sorting and analysed single unit activity, A.D. performed all immunohistochemistry and image analysis, H.H. provided resources for the experimental work and supervised the project.

H.H. conceptualized and coordinated the project. The manuscript was written by H.H. with significant contributions from all authors.

