## Supplementary Material for "Early-life stress impairs development of functional interactions and neuronal activity within prefrontal-amygdala networks *in vivo*"

Figure S1.

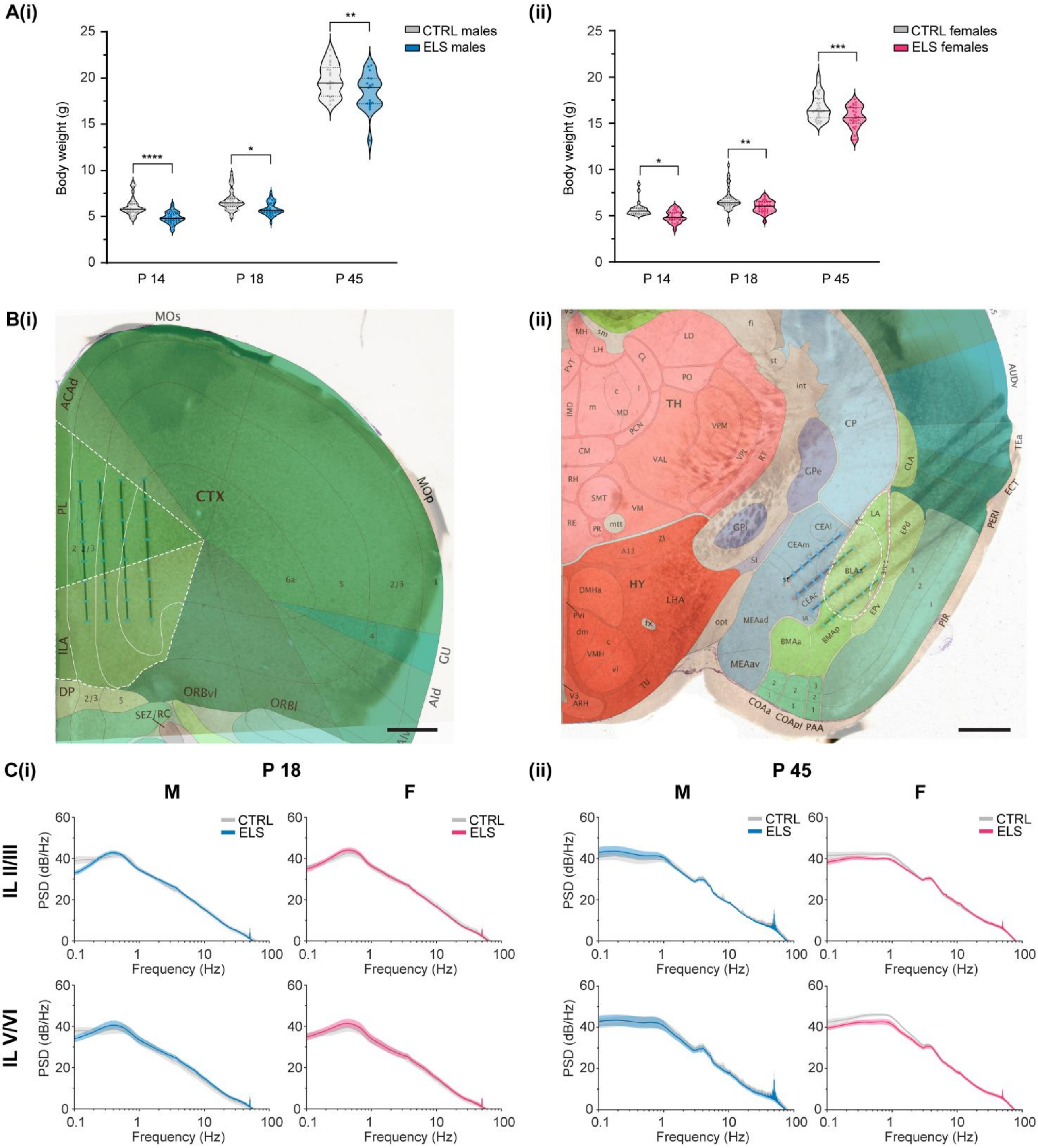


**Figure S1. Effect of ELS on body weight and oscillatory network activity within pre-juvenile and adolescent infralimbic networks. A, (i)** Violin plots showing body weight of control (grey) or ELS (blue) male mice at P14 immediately after the LBN paradigm, at P18 during pre-juvenile development, and at P45 during adolescence. Note the significantly lower body weight of ELS pups vs controls at all developmental time points. **(ii)** Same as A (i) for female control (grey) and ELS (pink) pups. See Suppl. Table 1 for the exact number of pups per group in each graph. **B,** Digital photomontage of a sample brightfield image with electrode tracts and reconstruction of the 32 recording channels (blue dots). The corresponding plate of the Allen Brain Atlas is superimposed, showing the electrode locations within **(i)** PL and IL or **(ii)** BLAa. Scale bar: 500 µm. **C, (i)** Average logarithmic power spectra of infralimbic LFP in layers II/III (top) or layers V/VI (bottom) of male (left)

and female (right) control (grey) or ELS (blue or magenta) pre-juvenile mice (males: N = 16-20/group, females: N = 12-18 /group). Note the broad peak in delta range (0.3-0.5 Hz) corresponding to slow-wave activity as well as the small peak in low-theta range at 4 Hz. **(ii)** Same as B (i) for adolescent control and ELS male and female mice (males: N = 13-15/group, females: N = 15-21/group). Note the shift of the peak in delta activity towards 1 Hz and the prominent broad peak in low-theta activity (3-5 Hz). See Suppl. Table 1 for the exact number of pups per group in each graph.

Figure S2.

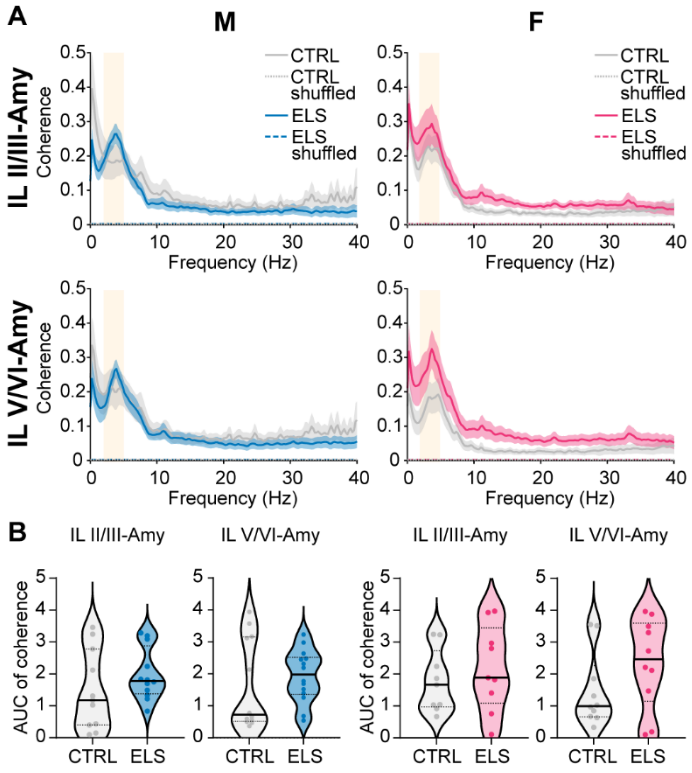


**Figure S2. ELS does not affect oscillatory coupling within infralimbic-amygdala networks in pre-juvenile mice. A,** Average coherence spectra of LFP in IL, layers II/III and the LFP in BLAa (top), or LFP in IL, layers V/VI and the LFP in BLAa (bottom) of male (left) and female (right) control (grey) or ELS (blue or magenta) pre-juvenile mice (males: N = 11-14/group, females: N = 9-10 /group). Note the broad peak in coherence in the low-theta range (3-5 Hz) in all groups, which is similar in control and ELS male and female mice. **B,** Violin plots with median and IQR showing the area under the curve (AUC) for the coherence in the low-theta band (3-5 Hz) within infralimbic layer II/III – BLAa or infralimbic layer V/VI – BLAa networks in male (left) and female (right) control (grey) or ELS (blue or magenta) pre-juvenile mice. Note that low-theta coherence within infralimbic layer II/III – BLAa networks is similar in pre-juvenile control and ELS mice for both males and females. See Suppl. Table 3 for the exact number of pups per group in each graph.

Figure S3.

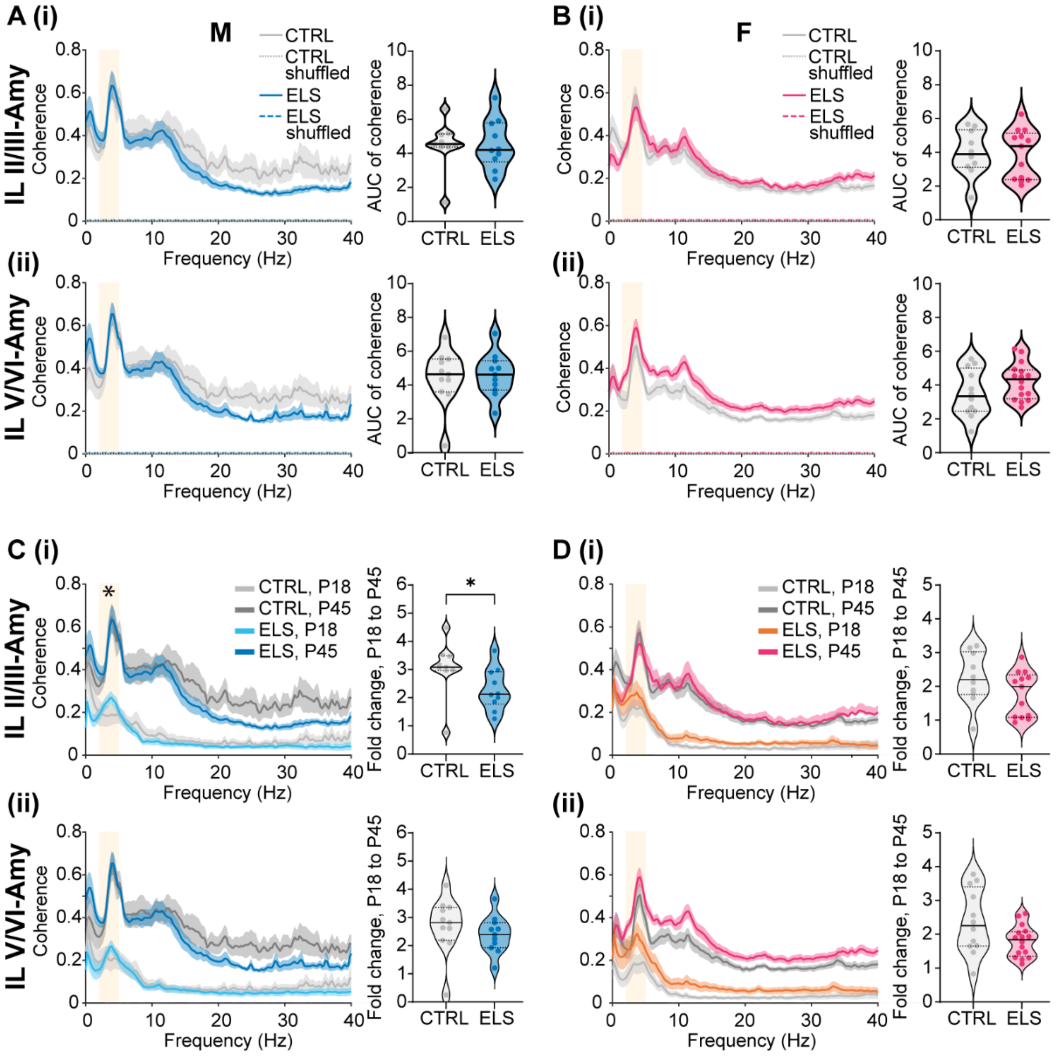


**Figure S3. ELS does not affect oscillatory coupling within adolescent infralimbic-amygdala networks. A, (i)** Average coherence spectra of the LFP in IL, layers II/III and the LFP in BLAa, and corresponding violin plots with median and IQR showing the area under the curve (AUC) for coherence in the low-theta band (3-5 Hz) in adolescent control (grey, N = 9) and ELS (blue, N = 11) male mice. **(ii)** Same as A (i) for coherence between the LFP in IL, layers V/VI and the BLAa (control: N = 10, ELS: N = 11). **B, (i)** Same as A (i) for adolescent control (grey, N = 10) and ELS (magenta, N = 12) female mice. **(ii)** Same as A (ii) for adolescent control (grey, N = 13) and ELS (magenta, N = 16) female mice. **C, (i)**, Average coherence spectra between the LFP in IL, layers II/III and the LFP in the BLAa for control pre-juvenile (light grey) and adolescent (dark grey) as well as ELS pre-juvenile (cyan) and adolescent (blue) male mice. The corresponding violin plots with median and IQR (right) show the fold change in low-theta coherence (3-5 Hz) between pre-juvenile and adolescent control (grey) or pre-juvenile and adolescent ELS (blue) male mice. Note that the fold increase in low-theta coherence is significantly smaller in ELS mice, suggesting that ELS promotes precocious development of functional interactions within these networks. **(ii)** Same as C (i) for coherence between the LFP in IL, layers V/VI and the LFP in the BLAa. **D, (i)** Same as C (i) for female pre-juvenile (light grey) and adolescent (dark grey) control and pre-juvenile (orange) and adolescent (magenta) ELS mice. The corresponding violin plots with median and IQR (right) show the fold change in low-theta coherence (3-5 Hz) between pre-juvenile and adolescent control (grey) or pre-juvenile and adolescent ELS (magenta) female mice. **(ii)** Same as D (i) for coherence between the

LFP in IL, layers V/VI and the LFP in the BLAa. See Suppl. Table 3 for the exact number of pups per group in each graph.

Figure S4.

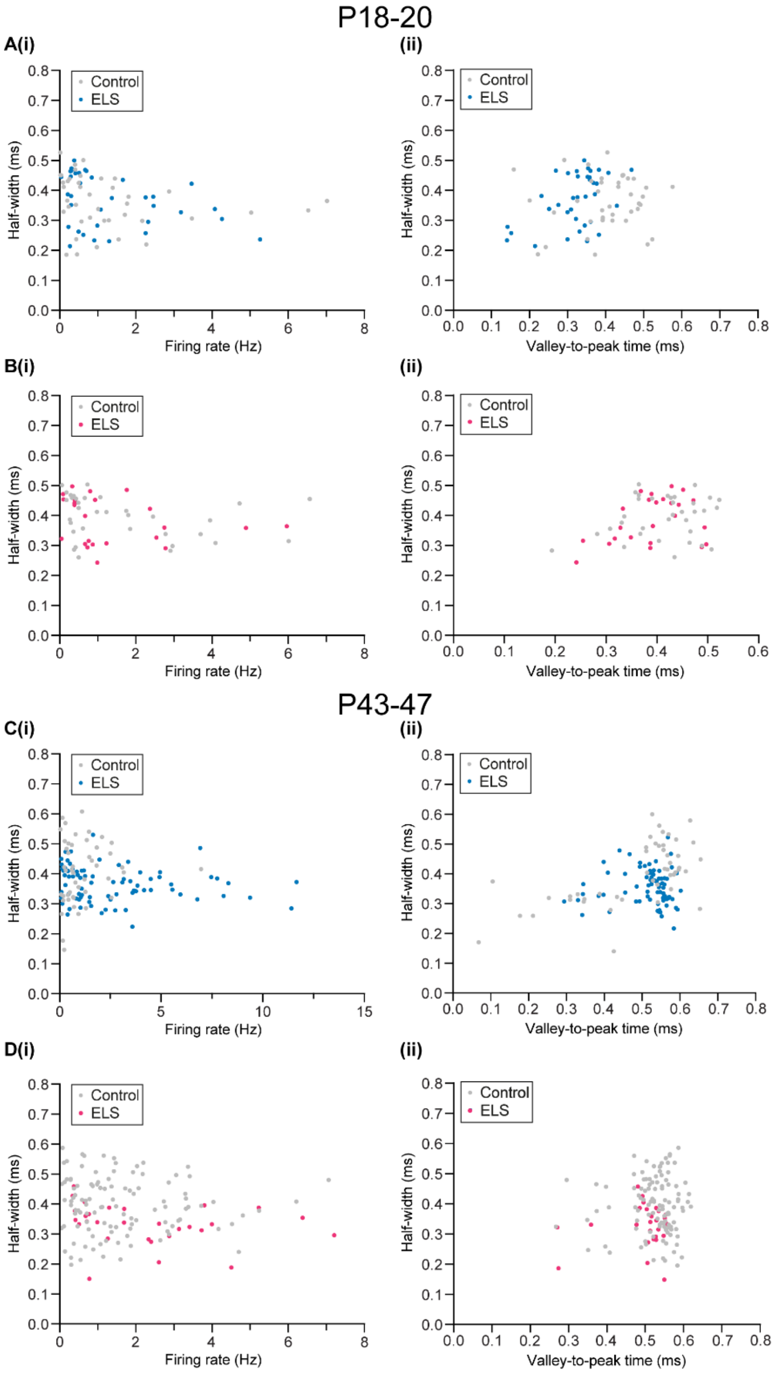


**Figure S4. Clustering of recorded BLAa units does not reveal association between firing rate and half-width and suggests that recorded neurons were mostly principal neurons in both groups in all experimental groups. A, (i)** Scatter plot showing firing rate as a function of half-width of the action potential waveform for all single units recorded from BLAa in pre-juvenile male control (grey) or ELS (blue) mice. Note that there is no association between firing rate and half-width. **(ii)** Scatter plot showing valley-to-peak time as a function of the half-width of the action potential waveform for all single units recorded from BLAa in pre-juvenile male control (grey) or ELS (blue) mice. Note that only a single cluster emerges for each group which overlap. **B, (i)** Same as A (i) for pre-juvenile female control (grey) and ELS (magenta) mice. **(ii)** Same as A (ii) for pre-juvenile female control (grey) and ELS (magenta) mice. **C, (i)** Same as A (i) for adolescent male control (grey) and ELS (blue) mice. **(ii)** Same as A (ii) for adolescent male control (grey) and ELS (blue mice. **D, (i)**

Same as B (i) for adolescent female control (grey) and ELS (magenta) mice. **(ii)** Same as B (ii) for adolescent female control (grey) and ELS (magenta) mice.

Figure S5.

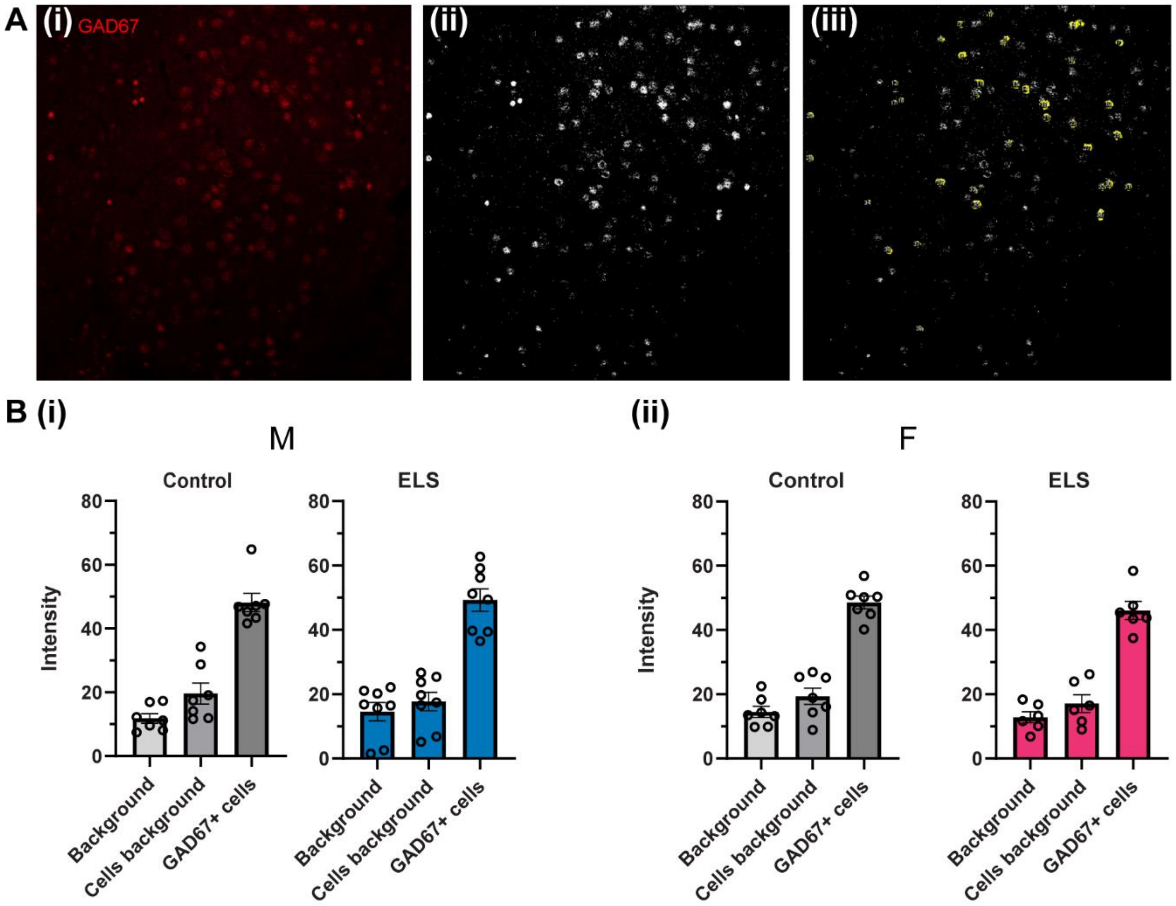


**Figure S5. Signal intensity of GAD67$^+$ BLA neurons vs background. A, (i)** Example microscope image of GAD67 immunofluorescent staining within the BLA of a P18 male control mouse. **(ii)** Same image as in (i) after thresholding in ImageJ. **(iii)** Same image as in (i) and (ii) with yellow markings indicating the cells identified by ImageJ for analysis. **B,** Intensity of the fluorescent signal as emitted from background (tissue without visible cells vs cells within background) versus signal from GAD67$^+$ neurons of (i) control and ELS P18 male mice or (ii) control and ELS P18 female mice. Individual data points correspond to individual brain sections analysed in each group. Note that the signal intensity of GAD67$^+$ cells was clearly distinct and much higher in all experimental groups compared to the signal from background.

Figure S6.

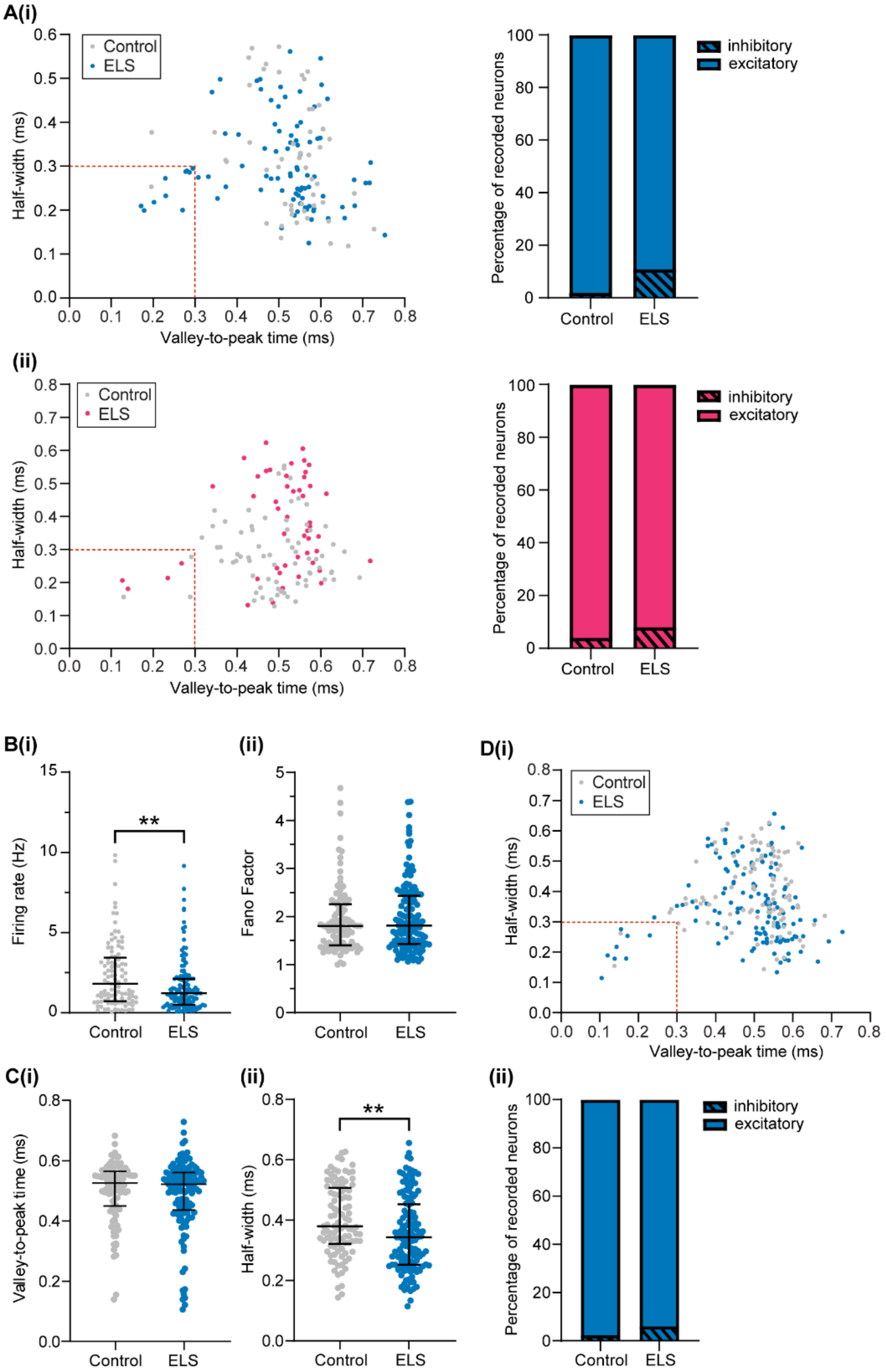


**Figure S6. ELS also decreases neuronal firing activity in IL networks. A, (i)** Left, scatter plot showing valley-to-peak time as a function of the half-width of the action potential waveform for all the single units recorded from PL layer II/III in pre-juvenile male control (grey) or ELS (blue) mice. Red dotted lines display the classification threshold for putative excitatory and putative inhibitory units (outside and inside the dotted square, respectively). Right, percentage of putative excitatory vs inhibitory neurons in the PL in each treatment group. **(ii)** Same as (i) for pre-juvenile female control (grey) and ELS (magenta) mice. **B, (i)** Distribution and median (IQR) of instant firing rates (Hz) of single units in IL layer V/VI in control (grey, n = 101 units from N = 13 pups) and ELS (blue, n = 137 units from N = 18 pups) male pre-juvenile mice. Note the significant decrease in firing rates after

ELS. **(ii)**, Same as B (i) for Fano factor. **C, (i)** Distribution and median (IQR) of the mean valley-to-peak time (ms) corresponding to the mAHP of the extracellular action potential of single units in the IL in control (grey, n = 101 units) and ELS (blue, n = 137 units) male pre-juvenile mice. **(ii)** Same as C (i) for the full width at half maximum (ms) of the extracellular action potential waveform. Note that the action potential half-width is decreased in ELS mice. **D, (i)** Same as A (i) for all single units recorded from IL layer V/VI in pre-juvenile male control (grey) or ELS (blue) mice. **(ii)** Percentage of putative excitatory vs putative inhibitory neurons in the IL in each treatment group. **$p < 0.01$, Mann-Whitney test.

Figure S7.

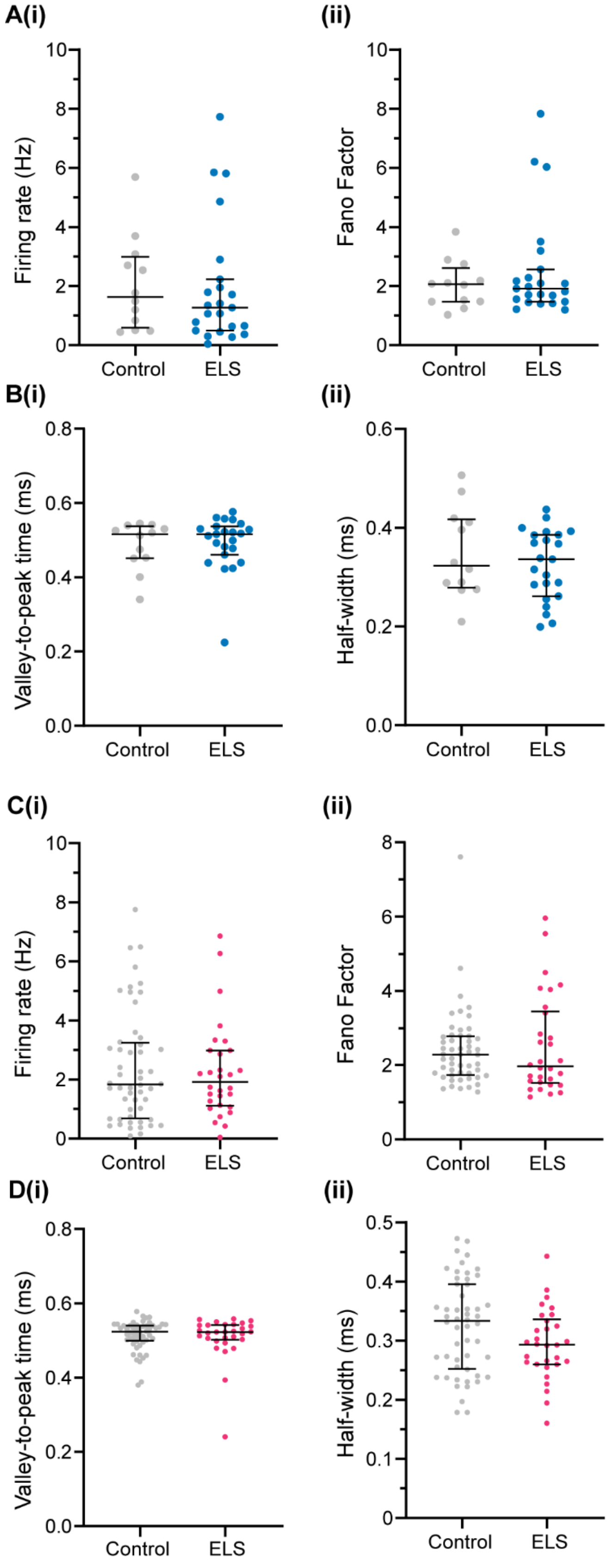
A(i)
(ii)
Firing rate (Hz)
Fano Factor
Control
ELS
B(i)
(ii)
Valley-to-peak time (ms)
Half-width (ms)
C(i)
(ii)
D(i)
(ii)

**Figure S7. ELS does not affect the firing activity nor the action potential parameters of neurons in layers II/III of the prelimbic cortex in adolescent male and female mice. A, (i)** Distribution and median (IQR) of instant firing rates (Hz) of single units in PL in control (grey, n = 12 units from N = 6 pups) and ELS (blue, n = 23 units from N = 9 pups) male adolescent mice. **(ii)**, Same as A (i) for Fano factor. **B, (i)** Distribution and median (IQR) of the mean valley-to-peak time (ms) corresponding to the mAHP of the extracellular action potential of single units in the PL in control (grey, n = 12 units) and ELS (blue, n = 23 units) male adolescent mice. **(ii)** Same as B (i) for the full width at half maximum (ms) of the extracellular action potential waveform. **C, (i)** Distribution and median (IQR) of instant firing rates (Hz) of single units in PL in control (grey, n = 52 units from N = 14 pups) and ELS (magenta, n = 30 units from N = 14 pups) female adolescent mice. **(ii)**, Same as C (i) for Fano factor. **D, (i)** Distribution and median (IQR) of the mean valley-to-peak time (ms) of single units in the PL in control (grey, n = 52 units) and ELS (magenta, n = 30 units) female adolescent mice. **(ii)** Same as D (i) for the full width at half maximum (ms) of the extracellular action potential waveform.

Figure S8.

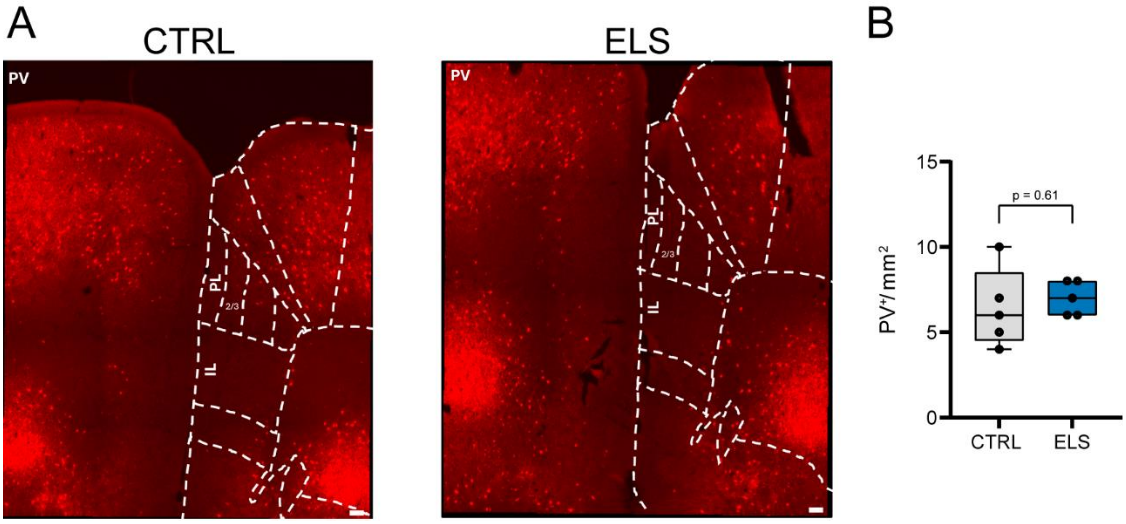


**Figure S8. Immunolabelling and cell density of parvalbumin-immunopositive (PV+) interneurons in prelimbic cortex (PL) of control (N=5) and ELS (N=5) P18 male mice. A,** Example microscope images of PV immunofluorescent staining within mPFC of a P18 male control (left) and P18 male ELS mouse (right). **B**, Box and whisker plots of cell density of PV+ neurons in PL in P18 control (N=5, grey) and ELS (N=5, blue) male mice. Note that PV+-cell density is not significantly different between the groups.

| | ***male*** | | ***female*** | |
|---|---|---|---|---|
| | **control** | **ELS** | **control** | **ELS** |
| *P14* | 32 | 38 | 26 | 24 |
| *P18-20* | 46 | 40 | 45 | 31 |
| *P43-47* | 21 | 18 | 28 | 26 |

**Suppl. Table 1**. Number of pups included in the calculation of the weight in Suppl. Figure 1 for all groups on the last day of the LBN protocol (P14), during pre-juvenile (P18), or adolescent (P45) development.

| | P 18 | | | | P45 | | | |
|---|---|---|---|---|---|---|---|---|
| | **male** | | **female** | | **male** | | **female** | |
| | **control** | **ELS** | **control** | **ELS** | **control** | **ELS** | **control** | **ELS** |
| *PL II/III* | 18 | 16 | 18 | 13 | 13 | 15 | 18 | 20 |
| *PL V/VI* | 20 | 20 | 18 | 14 | 16 | 16 | 21 | 22 |
| *IL II/III* | 17 | 16 | 16 | 12 | 13 | 13 | 15 | 18 |
| *IL V/VI* | 18 | 20 | 18 | 14 | 15 | 14 | 18 | 21 |
| *Amy* | 13 | 14 | 12 | 12 | 11 | 13 | 13 | 17 |

**Suppl. Table 2**. Number of pups included in the calculation of power spectra in Figure 1 or Suppl. Figure 1 in either PL or IL for all groups during pre-juvenile (P18) or adolescent (P45) development.

| | P 18 | | | | P45 | | | |
|---|---|---|---|---|---|---|---|---|
| | male | | female | | male | | female | |
| | control | ELS | control | ELS | control | ELS | control | ELS |
| *PL II/III-BLAa* | 11 | 12 | 10 | 11 | 9 | 12 | 11 | 15 |
| *PL V/VI-BLAa* | 12 | 14 | 10 | 11 | 11 | 13 | 13 | 17 |
| *IL II/III-BLAa* | 11 | 12 | 9 | 9 | 9 | 10 | 10 | 13 |
| *IL V/VI- BLAa* | 12 | 14 | 10 | 10 | 11 | 11 | 12 | 16 |

**Suppl. Table 3**. Number of pups included in the calculation of coherence spectra in Figure 2, Figure 3, Suppl. Figure 2, or Suppl. Figure 3 between PL-BLAa or IL-BLAa networks for all groups during pre-juvenile (P18) or adolescent (P45) development.

| **MALES** | **Two-way ANOVA** | | **POST-HOC TEST** | |
|---|---|---|---|---|
| **ΔFosB$^+$/mm$^2$** | ***F-value*** | ***p-value*** | **Age** | ***p-value*** |
| Interaction | F (1, 16) = 0.05873 | 0.8116 | P14 | 0.3428 |
| Treatment | F (1, 16) = 4.757 | **0.0444** | P18 | 0.2006 |
| Age | F (1, 16) = 1.279 | 0.2748 | | |
| **ΔFosB Intensity** | | | | |
| Interaction | F (1, 15) = 1.455 | 0.2464 | P14 | 0.5806 |
| Treatment | F (1, 15) = 0.03619 | 0.8517 | P18 | 0.7195 |
| Age | F (1, 15) = 1.455 | 0.2464 | | |
| **ΔFosB$^+$/mm$^2$/hemisphere** | **Three-way ANOVA** | | | |
| | ***F-value*** | | ***p-value*** | |
| Hemisphere | F (1, 29) = 0.1257 | | 0.7255 | |
| Age | F (1, 29) = 0.8827 | | 0.3552 | |
| Treatment | F (1, 29) = 6.718 | | **0.0148** | |
| Hemisphere x Age | F (1, 29) = 0.1892 | | 0.6668 | |
| Hemisphere x Treatment | F (1, 29) = 0.1767 | | 0.6773 | |
| Age x Treatment | F (1, 29) = 0.002258 | | 0.9624 | |
| Hemisphere x Age x Treatment | F (1, 29) = 0.07923 | | 0.7803 | |
| **FEMALES** | **Two-way ANOVA** | | **POST-HOC TEST** | |
| **ΔFosB$^+$/mm$^2$** | ***F-value*** | ***p-value*** | **Age** | ***p-value*** |
| Interaction | F (1, 12) = 9.132 | **0.0106** | P14 | 0.9588 |
| Treatment | F (1, 12) = 6.992 | **0.0214** | P18 | **0.0031** |
| Age | F (1, 12) = 17,79 | **0.0012** | | |
| **ΔFosB Intensity** | | | | |
| Interaction | F (1, 12) = 0.3218 | 0.5810 | P14 | 0.7085 |
| Treatment | F (1, 12) = 0.2801 | 0.6063 | P18 | 0.9995 |
| Age | F (1, 12) = 0.3218 | 0.5810 | | |
| **ΔFosB$^+$/mm$^2$/hemisphere** | **Three-way ANOVA** | | | |
| | ***F-value*** | | ***p-value*** | |
| Hemisphere | F (1, 22) = 0.0005045 | | 0.9823 | |
| Age | F (1, 22) = 17.98 | | **0.0003** | |
| Treatment | F (1, 22) = 10.67 | | **0.0035** | |
| Hemisphere x Age | F (1, 22) = 0.04087 | | 0.8417 | |
| Hemisphere x Treatment | F (1, 22) = 0.2272 | | 0.6383 | |
| Age x Treatment | F (1, 22) = 10.54 | | **0.0037** | |
| Hemisphere x Age x Treatment | F (1, 22) = 0.6339 | | 0.4345 | |

**Suppl. Table 4.** Statistical results related to Figure 5.
